# Brain–Language Alignment During Naturalistic Reading and Its Disruption by Mind-Wandering

**DOI:** 10.64898/2026.08.14.744875

**Authors:** Haorui Sun, David C. Jangraw

**Affiliations:** Department of Electrical and Biomedical Engineering, University of Vermont

**Keywords:** EEG encoding models, naturalistic reading, mind-wandering, word embeddings, fixation-related potentials, brain–language alignment

## Abstract

Encoding models offer a principled framework for linking computational representations of language to neural activity, but most electroencephalography (EEG) evidence for brain–language alignment comes from tightly controlled, word-by-word reading paradigms. Whether such alignment is detectable during naturalistic reading, and how it is affected by lapses in attention, remains unclear. We addressed these questions using ROAMM, a multimodal dataset containing simultaneous EEG and eye-tracking recordings with time-resolved mind-wandering (MW) annotations from 44 participants reading naturalistic texts. Ridge regression encoding models were trained to predict fixation-aligned EEG spectral power and fixation-related potentials (FRPs) from five word-embedding models (GloVe, word2vec, BERT, GPT-2, and Llama 3). Using permutation testing with false discovery rate correction, we found statistically reliable brain–language alignment across both feature types, with contextual embeddings outperforming static embeddings. Spectral alignment was strongest in the alpha and low-beta bands over parietal electrodes, while FRP-based alignment peaked 200–300 ms after fixation onset over central and parietal-occipital regions. Leveraging ROAMM’s span-level MW annotations, we further show that brain–language alignment is systematically reduced during MW, an effect that was substantially larger for oscillatory (PSD) than for event-related (FRP) features. These findings demonstrate that modern language-model representations are reflected in EEG activity during naturalistic reading despite the modality’s inherent noise, and that fluctuations in attention constitute an underappreciated source of variability in brain–language encoding studies.

## Introduction

Understanding how language is represented, processed, and transformed into meaning in the human brain has remained a central goal of cognitive neuroscience for decades (Caplan, 1987; Friederici, 2011; Poeppel et al., 2012). From lesion studies to modern neuroimaging approaches, previous research has identified neural signatures associated with lexical access (Indefrey & Levelt, 2004), semantic processing (Binder et al., 2009; Huth et al., 2016; Kutas & Hillyard, 1980), and discourse comprehension (Ferstl et al., 2007; Maguire et al., 1999), revealing that language understanding emerges from the coordinated activity of distributed brain networks (Friederici, 2011). These findings have provided important insights into where and when language-related processes occur in the brain. However, they often offer only a high-level characterization of language function and provide limited information about the specific linguistic content represented by neural activity. For example, when a reader processes a particular word, phrase, or sentence, it remains unclear what aspects of its meaning are encoded in neural responses and how these representations are distributed across the brain. Although behavioral measures and traditional experimental paradigms have substantially advanced our understanding of language processing, directly linking neural signals to the underlying linguistic representations remains a fundamental challenge, particularly during naturalistic language comprehension.

Recent advances in natural language processing (NLP) have created new opportunities for addressing this challenge. Early computational models represented words using discrete symbolic features, such as one-hot encodings or manually defined linguistic categories (Bengio et al., 2003; Jurafsky & Martin, 2009). While these representations allowed words to be represented computationally, they could not capture graded semantic relationships or encode words within a continuous vector space. The development of distributed word representations, commonly known as word embeddings (Mikolov et al., 2013; Pennington et al., 2014; Turney & Pantel, 2010), transformed this landscape by enabling words to be represented as continuous vectors in a high-dimensional space. In these embedding spaces, semantically related words tend to occupy nearby regions, allowing linguistic relationships to be quantified mathematically. In the last decade, large language models (LLMs) have dramatically expanded the expressive power of these representations. LLMs generate contextualized representations that incorporate information from surrounding text, enabling them to capture semantic, syntactic, and discourse-level information (Devlin et al., 2019; Peters et al., 2018). As a result, modern language models provide some of the richest computational representations of language currently available.

With these advances, two major approaches have emerged for investigating the relationship between linguistic representations and neural activity: encoding models and decoding models (Holdgraf et al., 2017; Naselaris et al., 2011). Decoding approaches attempt to predict linguistic content from neural signals. Given brain activity recorded during language comprehension, a decoding model seeks to reconstruct the perceived or intended words, sentences, or semantic concepts. Recent advances in neural decoding have demonstrated impressive results, including the reconstruction of perceived speech from electrocorticography (ECoG) and intracranial EEG (iEEG) recordings (Anumanchipalli et al., 2019; Chen et al., 2024; Pasley et al., 2012; Willett et al., 2023), as well as above-chance decoding of text from noninvasive neural measurements such as magnetoencephalography (MEG) and EEG (Défossez et al., 2023; d’Ascoli et al., 2025; Liu et al., 2024; Wang & Ji, 2022; Wang et al., 2024). These findings provide compelling evidence that rich linguistic information is embedded within neural activity. However, decoding models are often implemented using complex deep neural network architectures that operate largely as black boxes (Lipton, 2016). Moreover, their outputs are typically evaluated using linguistic similarity metrics between predicted and ground-truth text (Wang & Ji, 2022). While such approaches hold tremendous promise for applications including brain-computer interfaces and neural prosthetics, they often provide limited insight into which neural features or brain regions contribute to the recovered linguistic representations and how specific aspects of language are encoded in the brain.

Encoding models provide a complementary framework that is particularly well suited for addressing neuroscientific questions (Holdgraf et al., 2017; Naselaris et al., 2011). Rather than predicting language from neural activity, encoding models predict neural responses from computational representations of language (Goldstein et al., 2022, 2024, 2025a; Hollenstein et al., 2019; Kumar et al., 2024). If a linguistic representation successfully predicts physiological activity, this suggests that information contained within the representation is also reflected in the neural signal. Importantly, encoding models can be applied independently to individual electrodes, frequency bands, or temporal components, allowing researchers to characterize where and when specific linguistic information is represented in the brain (Goldstein et al., 2024; Kumar et al., 2024; Sato & Mizuhara, 2018). Consequently, encoding approaches offer a more direct method for investigating brain–language relationships than global decoding metrics, as they enable the spatial and temporal organization of linguistic representations to be examined explicitly.

Encoding models have produced promising results across a variety of language tasks and neural recording modalities. Mitchell et al. (2008) first demonstrated that semantic features derived from word co-occurrence statistics could predict distributed fMRI activation patterns associated with word meanings. Subsequent studies extended this framework to naturalistic language comprehension, revealing that semantic information is represented across widespread cortical networks and organized into distributed semantic maps throughout the brain (Huth et al., 2012, 2016). Encoding models have also been applied to electrophysiological recordings, with studies showing that word embeddings can predict EEG responses (i.e., event-related potentials) during reading and capture neural activity associated with semantic processing (Sassenhagen & Fiebach, 2020; Sato & Mizuhara, 2018; Schwartz & Mitchell, 2019). More recently, ECoG studies have shown that the hierarchical representations learned by large language models closely align with the spatial and temporal organization of language processing in the cortex (Goldstein et al., 2025a,b). Collectively, these findings demonstrate that encoding models provide a powerful computational framework for characterizing how linguistic information is represented across multiple spatial and temporal scales.

Despite these advances, several limitations remain. First, many successful brain–language encoding studies have relied on fMRI and ECoG recordings. While both modalities provide high signal-to-noise ratio neural measurements, they come with important tradeoffs (Haufe et al., 2018). fMRI offers excellent spatial resolution but relatively poor temporal resolution because neural activity is measured indirectly through slow hemodynamic responses (Logothetis et al., 2001). ECoG, in contrast, provides both high spatial and temporal resolution, but its invasive nature restricts recordings to clinical populations and limits the availability of large-scale datasets (Buzsáki et al., 2012).

Furthermore, both fMRI and ECoG studies are expensive and resource-intensive to collect. EEG offers a complementary alternative: it is noninvasive, relatively inexpensive, portable, and capable of capturing neural dynamics at the millisecond timescale. However, EEG signals typically exhibit lower signal-to-noise ratios and poorer spatial resolution, making it unclear whether robust brain–language relationships can be reliably detected.

Existing EEG-based encoding studies have primarily relied on highly controlled experimental paradigms. For example, Sassenhagen & Fiebach (2020) examined isolated word reading, whereas Schwartz & Mitchell (2019) used the ZuCo dataset (Hollenstein et al., 2018), which consists of sentence-level reading tasks. Although such paradigms prioritize experimental control and signal quality, they do not fully reflect the complexity of natural language comprehension encountered in everyday reading. Consequently, it remains unclear whether brain–language alignment can be reliably observed during more naturalistic reading, where neural signals are noisier and linguistic context unfolds over longer timescales.

Another important limitation concerns attentional state. Mind wandering, defined as a shift of attention away from the external task toward internally generated thoughts (Giambra, 1989; Smallwood & Schooler, 2006, 2015), occurs frequently during reading and other language tasks (Killingsworth & Gilbert, 2010; Seli et al., 2018). A large body of research has shown that MW alters both physiological and neural activity, including eye-movement behavior, pupil dynamics, EEG spectral power, and event-related potentials (Kam et al., 2022; Mézière et al., 2025; Steindorf & Rummel, 2020). Moreover, MW reliably impairs reading comprehension and memory for textual information, suggesting that the processing of external linguistic input is reduced during periods of attentional disengagement (Ebbert et al., 2024; Farley et al., 2013; Mooneyham & Schooler, 2013; Smallwood et al., 2007, 2008; Unsworth & McMillan, 2013; Wammes et al., 2016). Despite these findings, previous brain–language encoding studies have largely ignored fluctuations in attention and implicitly assumed continuous engagement with linguistic stimuli (Goldstein et al., 2022, 2024, 2025a; Hollenstein et al., 2019; Kumar et al., 2024). To our knowledge, only one previous study has directly examined the effects of MW on a brain–language encoding task, and that study focused on auditory language processing rather than naturalistic reading (Chen et al., 2025). As a result, attentional state may represent an important but largely overlooked source of variability in estimates of brain–language alignment.

These limitations motivate two central questions addressed in this study. First, how reliably can brain–language alignment be detected from EEG recordings collected during naturalistic reading, despite the lower signal-to-noise ratio of the modality and the complexity of the reading environment? Second, if such alignment exists, how is it influenced by attentional state, particularly episodes of mind wandering?

To address these questions, we investigated the relationship between linguistic representations and neural activity during naturalistic reading using ROAMM (Sun et al., 2026b), a simultaneous EEG and eye-tracking dataset that includes time-resolved mind-wandering annotations. ROAMM combines EEG, eye tracking, and time-resolved MW labels, providing a unique opportunity to examine how attentional state influences brain–language alignment. We employed an encoding-model framework in which word embeddings derived from both static and contextual language models were used to predict fixation-aligned EEG responses. Because naturalistic reading involves substantial variability in both neural activity and linguistic content, and because EEG measurements are inherently noisy, we expected brain–language correspondence to be modest in magnitude. Nevertheless, we hypothesized that fixation-aligned EEG responses would exhibit statistically reliable alignment with the linguistic representations of fixated words.

We further hypothesized that brain–language alignment would be disrupted during mind wandering (Chen et al., 2025) based on the perceptual decoupling theory (Smallwood & Schooler, 2006), which proposes that attention becomes disengaged from external sensory input and redirected toward internally generated thought. During attentive reading, neural activity is expected to remain closely coupled to the processing of external linguistic input, resulting in stronger correspondence between linguistic representations and neural responses. In contrast, during MW, cognitive resources are redirected toward internally generated thoughts, reducing the extent to which ongoing neural activity reflects the linguistic properties of the text. Consequently, we predicted that encoding performance would be significantly weaker during MW than during normal reading. To test these hypotheses, we used encoding models to predict EEG spectral features from word embeddings derived from both static and contextual language models and examined how encoding performance varied across frequency bands, scalp regions, and attentional states.

This study investigates whether word embeddings from modern language models can predict neural activity during naturalistic reading and whether this relationship is modulated by attentional state. By combining naturalistic EEG recordings, computational language models, and time-resolved mind-wandering annotations, this work extends brain–language encoding research beyond highly controlled laboratory paradigms and provides new insights into how fluctuations in attention influence the correspondence between linguistic representations and neural activity.

## Materials and Methods

To quantify the relationship between linguistic representations and neural activity during reading, we employed an encoding-model framework in which word embeddings were used to predict fixation-aligned EEG features (Figure 1). Static and contextual embeddings were paired with fixation-level power spectral density (PSD) and fixation-related potential (FRP) measures. Embeddings were used as input to ridge regression models trained to predict individual EEG features. Model performance was evaluated using a five-fold page-level cross-validation procedure and quantified as the Pearson correlation between predicted and observed neural responses. To examine the effect of mind wandering on brain–language alignment, the same encoding framework was applied to a balanced subset containing both normal-reading and MW fixations, and model performance was compared between the two attentional states. These steps are described in detail in the following sections.

**Figure 1.**
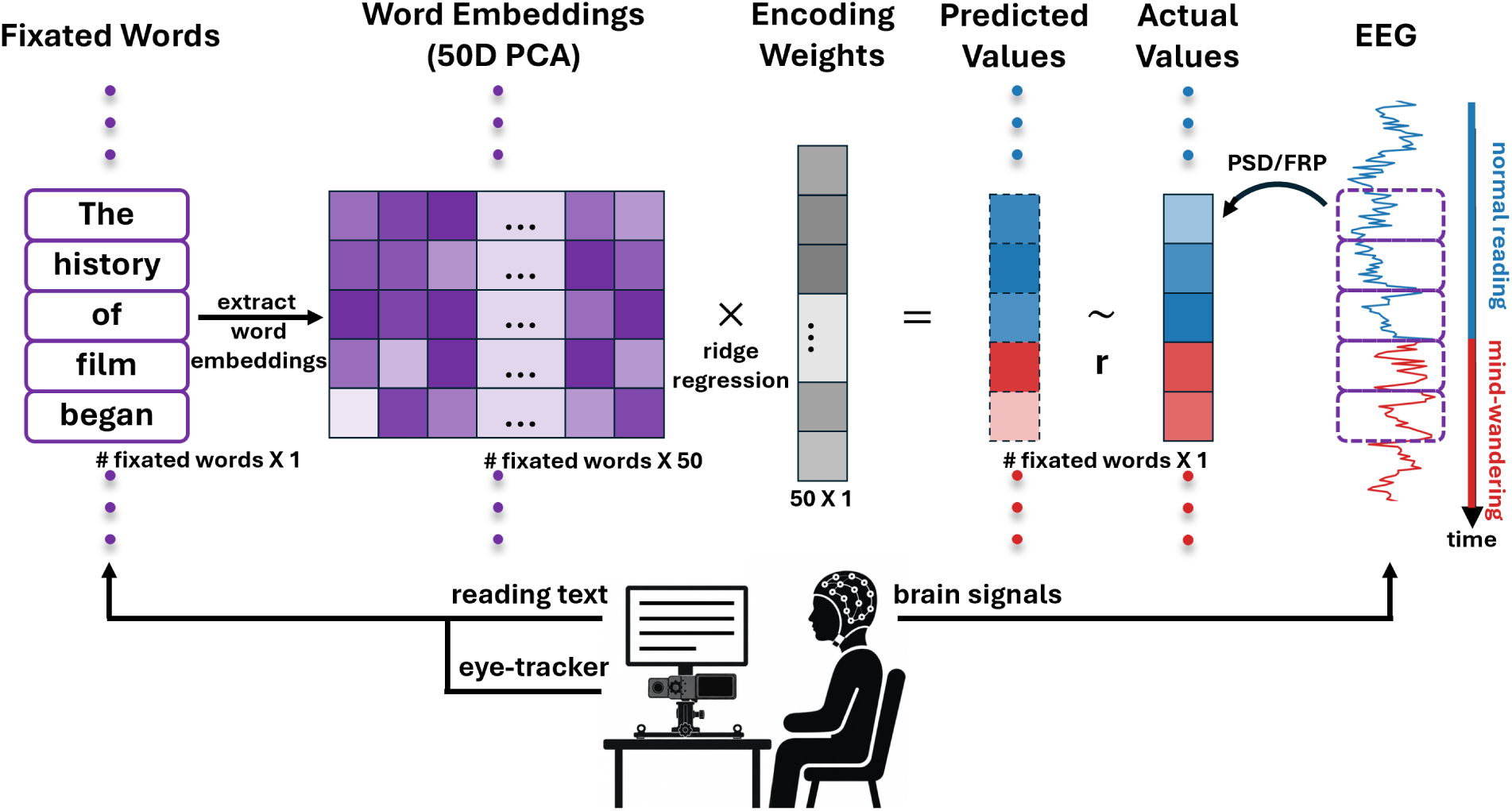
Encoding model framework. Five word embedding models (GloVe, word2vec, BERT, GPT-2, and Llama 3) were used to predict fixation-aligned EEG spectral power and fixation-related potential features. Embeddings were reduced using principal component analysis (PCA), and separate ridge regression models were trained for each EEG feature. Model performance was quantified as the correlation between predicted and observed neural responses.

### Data

Rather than collecting new data, this study reuses the ROAMM (Reading Observed at Mindless Moments) dataset (Sun et al., 2026b), a publicly available multimodal resource containing synchronized EEG and eye-tracking recordings collected while participants read naturalistic texts and self-reported episodes of mind wandering. All data collection, participant recruitment, and mind-wandering annotation were carried out as part of the original ROAMM data release and are described in full by Sun et al. (2026b); the aspects of that protocol most relevant to the present encoding analyses are summarized below for completeness.

As reported by Sun et al. (2026b), participants were recruited from a university community and screened for fluent English reading ability and no family history of neurological disorders or epilepsy, provided informed consent prior to participation, and were tested under a protocol approved by the data collectors’ Institutional Review Board. After excluding participants with incomplete recordings, monocular-only eye-tracking data, or missing demographic information, the released dataset retained a final sample of 44 participants (ages 18–64 years, median = 20, mean = 22.6; 30 female, 9 male, 5 non-binary; 37 right-handed, 7 left-handed).

During the original ROAMM data collection, participants read five Wikipedia articles (Pluto, The Prisoner’s Dilemma, Serena Williams, The History of Film, and The Voynich Manuscript), each divided into 10 pages of approximately 220 words displayed as plain text, and answered multiple-choice comprehension questions after each article.

Mind-wandering episodes were annotated using the ReMind retrospective self-report paradigm (Sun et al., 2026a): whenever participants noticed their attention had drifted from the text, they pressed a button and then retrospectively identified the words at which their mind had begun and stopped wandering. MW onset was defined as the time of the first fixation on the self-reported onset word, and MW offset was defined as 2 s before the report button press. Using this procedure, participants reported 1,001 MW episodes across approximately 45.5% of the 2,200 reading pages in the released dataset.

Binocular eye movements and pupil area were recorded at 1000 Hz using an EyeLink 1000 Plus system (SR Research), and simultaneous EEG was recorded from 64 channels using a BioSemi ActiveTwo system at a sampling rate of 2048 Hz. Eye-tracking and EEG data streams were synchronized using page-onset triggers, and EEG signals were resampled to 256 Hz, re-referenced, band-pass filtered (0.5–50 Hz), and corrected for bad channels and ocular and muscle artifacts using independent component analysis prior to release. In total, the released dataset comprises over 46 million samples from the 44 participants (more than 50 hours of synchronized recording), including approximately 26 million samples (about 30 hours) of first-pass reading and approximately 2.2 aggregate hours of annotated mind wandering, all of which were available for the encoding analyses reported here.

### Linguistic Representations

We evaluated five word embedding models that have been widely used in previous studies of brain–language alignment employing encoding-model frameworks (Goldstein et al., 2022, 2024, 2025a; Hollenstein et al., 2019). Static word embeddings were generated using the GloVe (Pennington et al., 2014) and word2vec (Mikolov et al., 2013) algorithms, which assign a single vector representation to each word regardless of context. Contextual embeddings were extracted from BERT Base Uncased (Devlin et al., 2019), GPT-2 Small (Radford et al., 2019), and Meta Llama 3 8B (Grattafiori et al., 2024). For the contextual models, embeddings were generated using overlapping sliding windows of up to 512 tokens with a 128-token overlap, and representations were extracted from the final hidden layer (Goldstein et al., 2024). When a word appeared in multiple windows, its final representation was computed by averaging the contextual embeddings across all occurrences. Stop words, defined as common words that primarily serve grammatical functions rather than conveying substantial semantic meaning (e.g., the, and, of), were excluded from all analyses using the English stop-word list provided by the Natural Language Toolkit (NLTK) (Bird, 2006).

Embeddings were aligned to fixation-level EEG data using fixation-to-word mappings within each story. Consequently, each fixation was associated with both a linguistic representation of the fixated word and a corresponding neural response. Prior to model fitting, embedding vectors were standardized using statistics derived from the training data within each cross-evaluation fold. To reduce dimensionality and mitigate collinearity among embedding features, principal component analysis (PCA) was applied to the standardized embeddings, and the first 50 principal components were retained, similar to a previous study (Goldstein et al., 2025a). To prevent information leakage, both the standardization and PCA transformations were fit exclusively on the training data and subsequently applied to the held-out test data using the training-derived parameters.

### Neural Signals

Encoding analyses were performed separately for spectral and event-related EEG measures. Both feature types were extracted relative to individual fixation events using the fixation timing information available in the ROAMM dataset (Sun et al., 2026b). A total of 592 fixations exceeding 800 ms in duration were excluded as outliers, representing 0.15% of the 393,344 recorded fixations. Although fixation durations during reading are typically substantially shorter than 800 ms (Rayner, 1998), mind-wandering episodes are often associated with longer fixations (Reichle et al., 2010; Sun et al., 2026a). We therefore adopted a relatively conservative threshold to retain naturally occurring fixation events while excluding extreme values likely attributable to artifacts or noise.

For spectral analyses, fixation-aligned power spectral density features were computed for eight frequency bands: low theta (4.0–6.0 Hz), high theta (6.5–8.0 Hz), low alpha (8.5–10.0 Hz), high alpha (10.5–13.0 Hz), low beta (13.5–18.0 Hz), high beta (18.5–30.0 Hz), low gamma (30.5–40.0 Hz), and high gamma (40.0–49.5 Hz). Band definitions were adapted from those used in the ZuCo dataset (Hollenstein et al., 2018) and were identical to those used by Sun et al. (2026a) in a prior mind-wandering classification analysis, facilitating comparisons across studies. Note that unlike the fixed 2-s windows used in that analysis, spectral features in the present study were computed within each fixation window, such that the temporal extent of the analysis was determined by the duration of the fixation itself. Continuous EEG signals were first band-pass filtered within each frequency range using a zero-phase fourth-order Butterworth filter implemented in second-order sections (SOS) form. The Hilbert transform was then applied to obtain the analytic signal, and instantaneous band power was computed as the squared magnitude of the analytic signal. Band-power values were log-transformed and averaged across each fixation window to obtain fixation-aligned spectral features. PSD values were computed for all 64 EEG channels, resulting in 512 features per fixation (64 channels × 8 frequency bands). Outliers were subsequently removed independently within each channel and frequency band using the median absolute deviation (MAD) method (Leys et al., 2013) with a threshold of 3.5.

To examine the influence of the aperiodic EEG background on encoding performance, we additionally evaluated encoding models using 1/f-corrected spectral features. To account for the aperiodic component of the EEG power spectrum, spectral features were normalized using channel- and run-specific baseline estimates derived from the SpecParam framework (Donoghue et al., 2020). For each participant, story, and electrode, Welch power spectra were computed from all first-pass reading periods, excluding the final two seconds of each page. SpecParam was then used to fit the aperiodic component of the spectrum within the 4–49.5 Hz range using a fixed aperiodic model. The fitted offset and exponent parameters were used to estimate the expected aperiodic power at the center frequency of each spectral band. Fixation-level log-power values were subsequently corrected for the aperiodic component by subtracting the estimated log-power of the 1/f background at each frequency, thereby reducing the influence of broadband spectral differences and emphasizing oscillatory activity above the aperiodic background.

For event-related analyses, fixation-related potentials were extracted from epochs spanning 500 ms before fixation onset to 500 ms after fixation onset. FRPs were baseline corrected by subtracting the mean signal during the 500-ms pre-fixation interval. To characterize fixation-related neural activity while reducing dimensionality, FRPs were averaged within consecutive 100-ms time bins spanning 0–500 ms relative to fixation onset.

This resulted in five temporal features for each of the 64 electrodes, yielding 320 electrode-by-time-bin features per fixation. The use of 100-ms bins was motivated by the temporal structure of canonical FRP components observed during reading, including the P1, N1, P2, N2, P300, and N400 (Degno & Liversedge, 2020). As demonstrated by Sun et al. (2026a) in FRP analyses of naturalistic reading, these components exhibited relatively distinct temporal distributions. Averaging neural activity within 100-ms windows therefore provided a compact representation of fixation-related responses while preserving sensitivity to the major FRP components. Outliers were identified and removed independently for each electrode and time bin using the MAD method (Leys et al., 2013) with a threshold of 3.5.

### Encoding Model

Encoding models were implemented using ridge regression. For each EEG feature, a separate model was trained to predict neural responses from the 50-dimensional PCA-reduced embedding representation. Ridge coefficients were estimated by minimizing

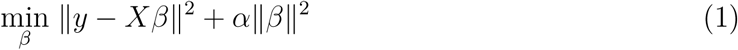

where X denotes the embedding features, y denotes a neural feature, and *α* = 10 is a fixed regularization parameter. This value was selected as a conservative regularization parameter to improve generalization and stabilize coefficient estimates when modeling high-dimensional embedding representations.

Neural targets were standardized using the mean and standard deviation of the training data within each fold. The resulting models generated predictions for held-out samples, which were evaluated in z-scored units relative to the training distribution.

### Model Evaluation

Encoding performance was quantified using the Pearson correlation coefficient between predicted and observed neural responses:

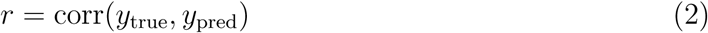

Although ridge regression was trained by minimizing mean squared error (MSE), prediction accuracy was evaluated using Pearson correlation. Correlation-based metrics are commonly used in neural encoding studies (Goldstein et al., 2022, 2024, 2025a; Hosseini et al., 2024) because they quantify the extent to which predicted responses capture fluctuations in the observed neural signal independent of overall signal magnitude or scaling. In contrast, MSE is sensitive to differences in amplitude and variance, making it less directly interpretable as a measure of representational correspondence. We therefore used Pearson correlation as our primary measure of brain–language alignment.

Model performance was evaluated using page-level five-fold cross-validation. Each story consisted of 10 pages, which were partitioned into five non-overlapping folds containing two pages each. For a given fold, data from the remaining eight pages served as the training set and data from the held-out pages served as the test set. All preprocessing operations, including feature standardization, PCA fitting, and target normalization, were performed independently within each training fold. Predictions from all held-out folds were concatenated to obtain a single out-of-sample prediction vector for each neural feature.

This cross-validation strategy was chosen to balance two competing goals. First, the encoding models should be trained on a sufficiently diverse set of linguistic contexts to learn meaningful relationships between word embeddings and neural responses. In particular, contextual language representations depend heavily on semantic and discourse information that may not generalize well when an entire story is withheld. Second, training and testing samples should remain temporally separated to reduce temporal dependencies between neighboring words and fixations, which could otherwise inflate performance estimates. Compared with word-level random train-test splits, the page-level approach reduces the likelihood of temporal leakage while preserving exposure to the semantic content of all stories. Compared with leave-one-story-out validation, it provides a less restrictive estimate of brain–language correspondence by allowing the model to learn from the full range of narrative contexts represented in the dataset. This approach was therefore selected as a compromise between semantic coverage and temporal independence.

Statistical significance was assessed using permutation testing. For each permutation, neural responses were randomly shuffled within story and page while preserving the distributional properties of the data. The complete encoding pipeline, including PCA fitting, model training, and evaluation, was repeated on the permuted data to generate a null distribution of prediction performance. Observed correlations were compared against the corresponding null distributions, and resulting p values were corrected for multiple comparisons using the false discovery rate (FDR) procedure (Benjamini & Hochberg, 1995).

### Model Weights and Semantic Interpretation

To examine whether different scalp regions relied on similar linguistic dimensions, we analyzed the similarity structure of the encoding model weights across EEG features. For each embedding model, neural feature type, signal component, and cross-validation fold, ridge-regression coefficients were extracted from the models. Pairwise Pearson correlations were computed between coefficient vectors across electrodes, yielding a 64 × 64 channel-by-channel weight similarity matrix. These matrices were then averaged across the five page-level cross-validation folds to obtain an estimate of the similarity structure for each embedding model and signal component. Specifically, for electrodes i and j, weight similarity was computed as

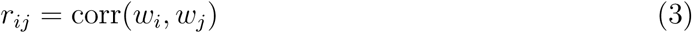

where w_i and w_j denote the ridge-regression coefficient vectors associated with the two electrodes.

To examine the extent to which the observed weight-similarity structure reflected the input linguistic representations or the output neural features, the same analysis was additionally applied to models trained using shuffled Llama 3 embeddings and to the corresponding raw neural features. This allowed the inter-electrode correlation structure of the recovered model weights to be compared with both the embedding input and the neural output.

To facilitate semantic interpretation of the learned weights, model coefficients were projected back into the original embedding space. Let w_PCA denote the ridge-regression coefficients learned in the PCA-transformed embedding space and P denote the corresponding PCA loading matrix. The recovered weight vector in the original embedding space was computed as

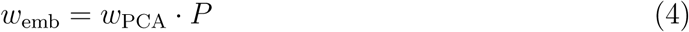

Each word embedding x_k was then projected onto the recovered weight vector,

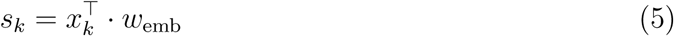

where s_k represents the contribution of word k to the predicted neural response. Words were ranked according to these scores, and the highest- and lowest-scoring words were identified as the top- and bottom-scoring words, respectively.

### Mind-Wandering Analysis

To examine whether MW influences brain–language alignment, we leveraged the span-level MW annotations available in the ROAMM dataset (Sun et al., 2026b). Fixations occurring within annotated MW spans were labeled as MW, whereas all remaining first-pass reading fixations were labeled as normal reading.

To ensure comparable sample sizes across attentional states, a balanced dataset containing equal numbers of normal-reading and MW fixations was constructed via random sampling without replacement. The same preprocessing, dimensionality reduction, and encoding procedures described above were then applied to this subset. For each embedding model and EEG feature, encoding models were trained using the balanced training data and evaluated separately on held-out normal-reading and MW test samples.

Brain–language alignment was quantified as the Pearson correlation between predicted and observed neural responses. Encoding performance was then compared between normal-reading and MW conditions for each EEG feature. A reduction in encoding performance during MW was interpreted as evidence of weaker correspondence between linguistic representations and neural activity during periods of attentional disengagement. Statistical significance was assessed using permutation testing and corrected for multiple comparisons using the FDR procedure.

## Results

In this study, we investigated whether linguistic representations derived from modern language models are reflected in neural activity during naturalistic reading. Using an encoding-model framework, we evaluated the extent to which fixation-aligned EEG spectral power and fixation-related potentials could be predicted from static and contextual word embeddings. In this section, we first characterize the overall strength and spatial distribution of brain–language alignment across embedding models and neural signal types. We then examine how this relationship is modulated by attentional state by comparing encoding performance during normal reading and mind wandering. Together, these analyses provide insight into how linguistic information is represented in neural activity during natural reading and the extent to which this representation is disrupted when attention disengages.

### Overall Brain–Language Alignment

We first examined whether linguistic representations could predict fixation-aligned neural activity using the encoding framework described above. Encoding performance was quantified as the Pearson correlation between predicted and observed neural responses. For visualization, we summarized the distribution of significant encoding results (p < 0.01, FDR corrected) across embedding models using boxplots (Figure 2) and reported the number of neural features that exhibited significant brain–language alignment. For PSD analyses, the total number of possible targets was 512 (64 channels × 8 frequency bands), whereas FRP analyses included 320 targets (64 channels × 5 temporal windows).

**Figure 2.**
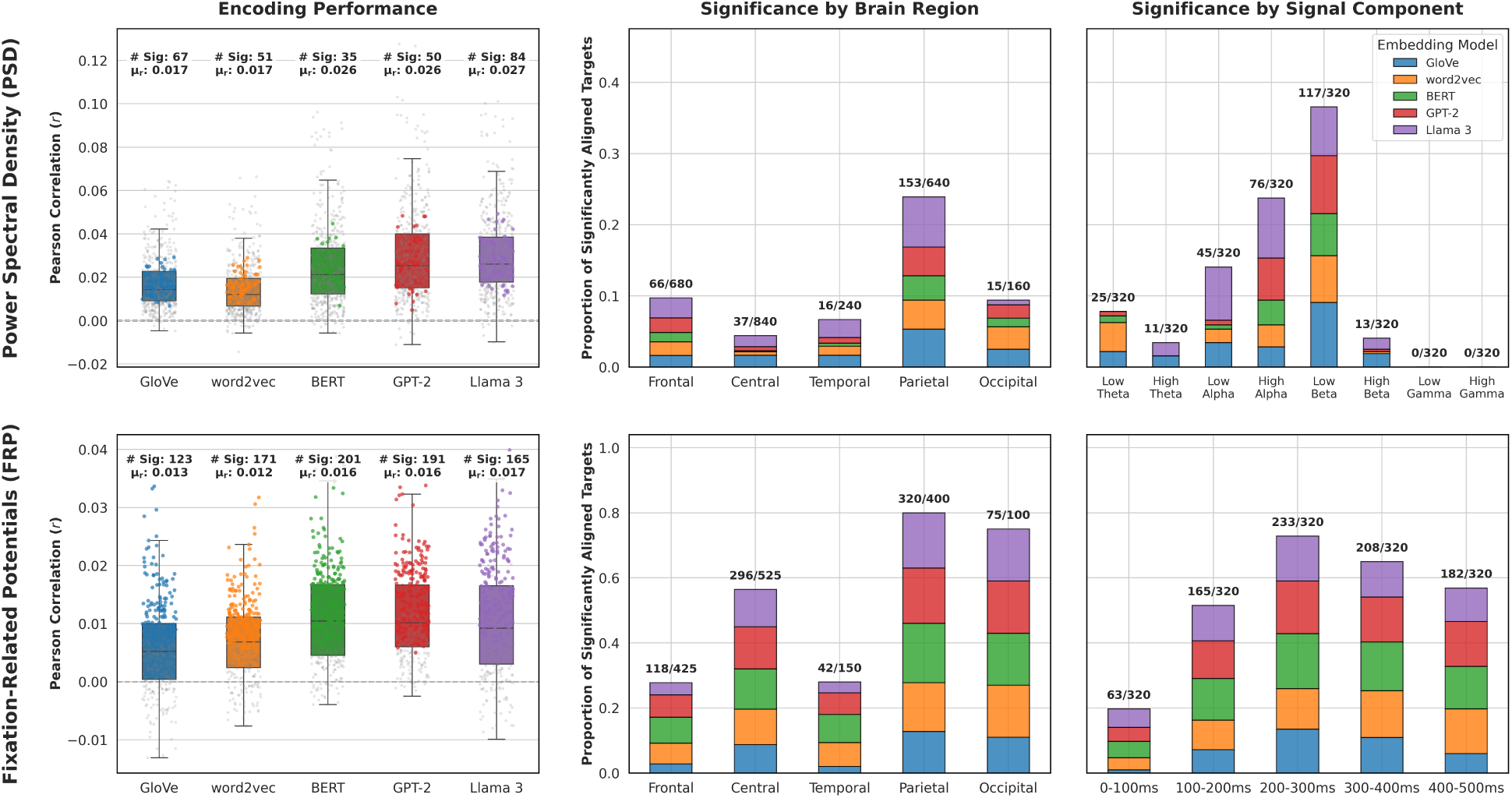
Brain–language alignment across embedding models for fixation-aligned EEG spectral power (PSD; top row) and fixation-related potentials (FRPs; bottom row). Left panels summarize the distribution of significant encoding correlations (p < 0.01, FDR corrected) across embedding models using boxplots. Numbers above each boxplot indicate the number of significantly aligned neural features and the mean correlation coefficient. Middle panels summarize the proportion of significantly aligned targets grouped by scalp region. Right panels summarize the proportion of significantly aligned targets grouped by signal component (frequency band for PSD; temporal window for FRPs).

We observed that contextual language models generally outperformed static embeddings. Among the significant encoding targets, contextual embeddings produced significantly higher encoding correlations than static embeddings for both PSD features (mean r = 0.0265 vs. 0.0172, Mann–Whitney p < 0.001) and FRP features (mean r = 0.0160 vs. 0.0125, Mann–Whitney p < 0.001). Among all embedding models, Llama 3 produced the largest number of significantly predicted PSD features, with 84 of 512 channel-by-frequency targets exhibiting significant brain–language alignment, whereas BERT produced the largest number of significant FRP features, with 201 of 320 channel-by-time-window targets reaching significance. Static embeddings also yielded significant encoding effects, although the number of significant targets was generally smaller and the corresponding correlation coefficients were weaker. When comparing PSD and FRP features, a substantially larger proportion of FRP features exhibited significant encoding effects.

### Spatial and Spectral Distribution of Encoding Performance

To characterize the distribution of significant encoding results, we computed the proportion of neural features exhibiting significant brain–language alignment across scalp regions and signal components (frequency bands for PSD and temporal windows for FRPs). Spatially, PSD encoding performance was most prominent over parietal electrodes, whereas FRP encoding effects were distributed more broadly across central, parietal, and occipital regions (Figure 2, middle column).

For PSD features, significant encoding effects were concentrated in the alpha and beta frequency ranges. Across all embedding models, low beta exhibited the highest proportion of significant targets (36.6%), followed by high alpha (23.8%) and low alpha (14.1%). In contrast, no significant effects were observed in the gamma bands (Figure 2, top right).

For FRP features, significant encoding effects were observed across most post-fixation time windows. Performance was weakest immediately following fixation onset (0–100 ms; 19.7% significant targets), and then increased substantially in later windows, reaching its peak during the 200–300 ms interval (72.8% significant targets), followed by the 300–400 ms (65%) and 400–500 ms (56.9%) windows (Figure 2, bottom right).

### Topographical Patterns of Brain–Language Alignment

To visualize the spatial distribution of encoding performance, we generated topographic maps showing the correlations between predicted and observed neural responses (Figure 3). For visualization purposes, we selected one representative static embedding model (GloVe) and one representative contextual embedding model (Llama 3), as these produced the strongest encoding effects among their respective categories. For PSD features, we present results for the alpha and beta bands, which exhibited the most robust brain–language alignment. For FRP features, we present the 0–100 ms, 200–300 ms, and 400–500 ms time windows to illustrate how encoding performance varied across stages of processing following fixation onset. Complete topographic results for all frequency bands and temporal windows are provided in the Supplementary Materials.

**Figure 3.**
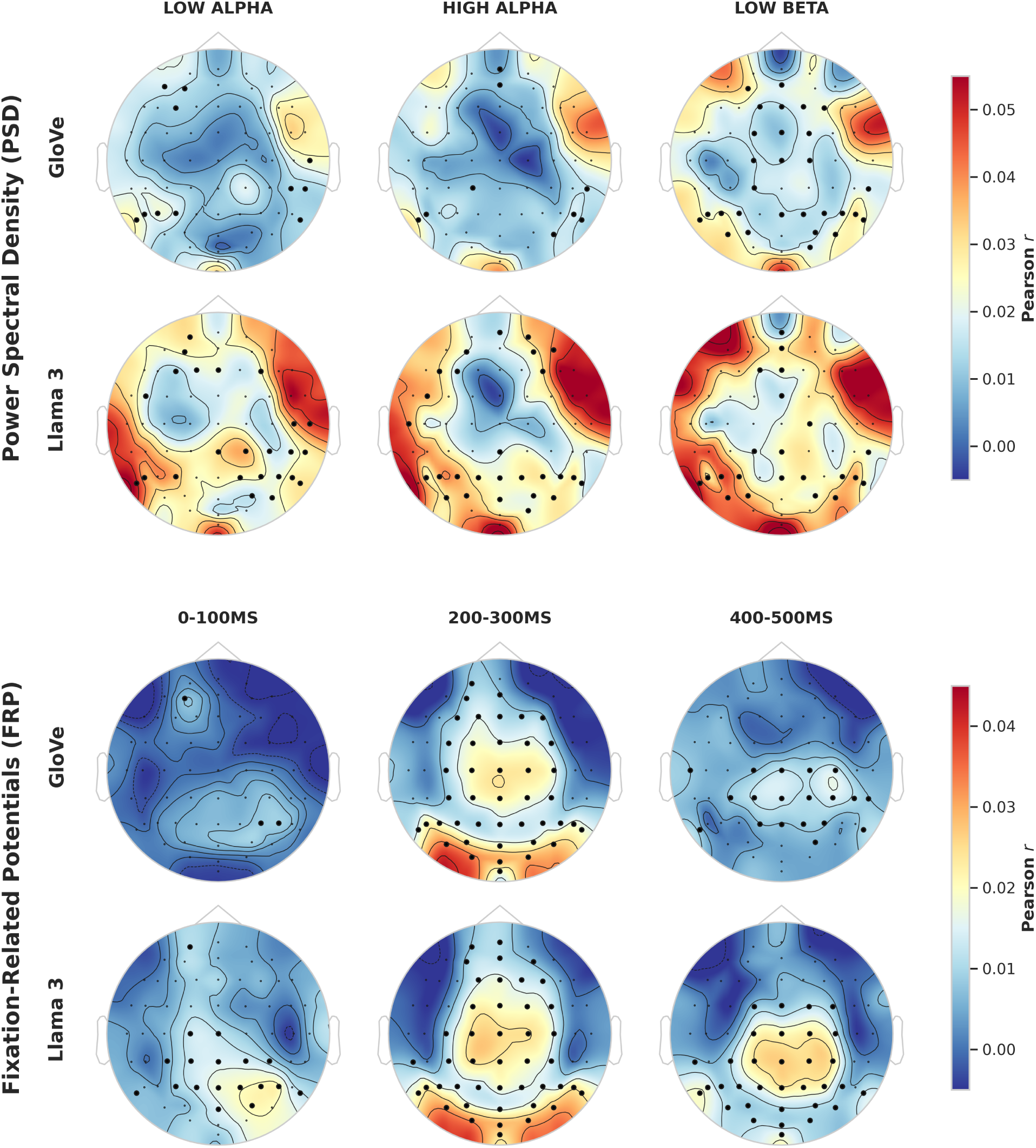
Topographical distribution of brain–language alignment for representative PSD frequency bands and FRP time windows. Color indicates the Pearson correlation between predicted and observed neural responses, averaged across electrodes. Black dots denote electrodes exhibiting significant encoding performance (p < 0.01, FDR corrected).

For PSD features, the strongest effects were observed in the low-beta and alpha frequency bands. In the low-beta band, both GloVe and Llama 3 exhibited significant encoding performance over central-frontal and temporo-parietal regions. Similar spatial patterns were observed in the alpha bands, although Llama 3 produced a larger number of significant electrodes than GloVe. These findings are consistent with the overall observation that contextual embeddings yielded stronger brain–language alignment than static embeddings.

Results obtained using 1/f-corrected PSD features were largely similar to those observed for the original PSD features (Supplementary Materials). The uncorrected PSD features exhibited numerically higher encoding correlations over peripheral electrodes, although these effects rarely reached statistical significance. Consequently, the spatial distribution and number of significant encoding results were largely comparable between the corrected and uncorrected analyses. In both cases, encoding effects in the higher frequency bands generally failed to survive permutation testing and FDR correction.

For FRP features, little significant encoding was observed immediately following fixation onset (0–100 ms). In contrast, the 200–300 ms window exhibited widespread significant effects over central, parietal, and occipital regions. These effects were present for both GloVe and Llama 3 embeddings, although the magnitude and spatial extent of the correlations were generally greater for Llama 3. Encoding performance gradually decreased during the 300–500 ms interval, although significant effects remained evident over centro-parietal regions, particularly for contextual embeddings.

### Spatial Structure of Encoding Model Weights

To examine the spatial organization of the recovered encoding-model weights, we computed inter-electrode correlation matrices for each neural feature. For visualization purposes, we selected representative results from GloVe and Llama 3 for high-alpha PSD and 200–300 ms FRP features (Figure 4). We additionally included shuffled Llama 3 embeddings and the corresponding raw neural features. Electrodes were grouped into cortical regions (frontal, central, temporal, parietal, and occipital) and ordered within each region according to their left (L), midline (M), and right (R) locations. Complete correlation matrices for all frequency bands and temporal windows are provided in the Supplementary Materials.

**Figure 4.**
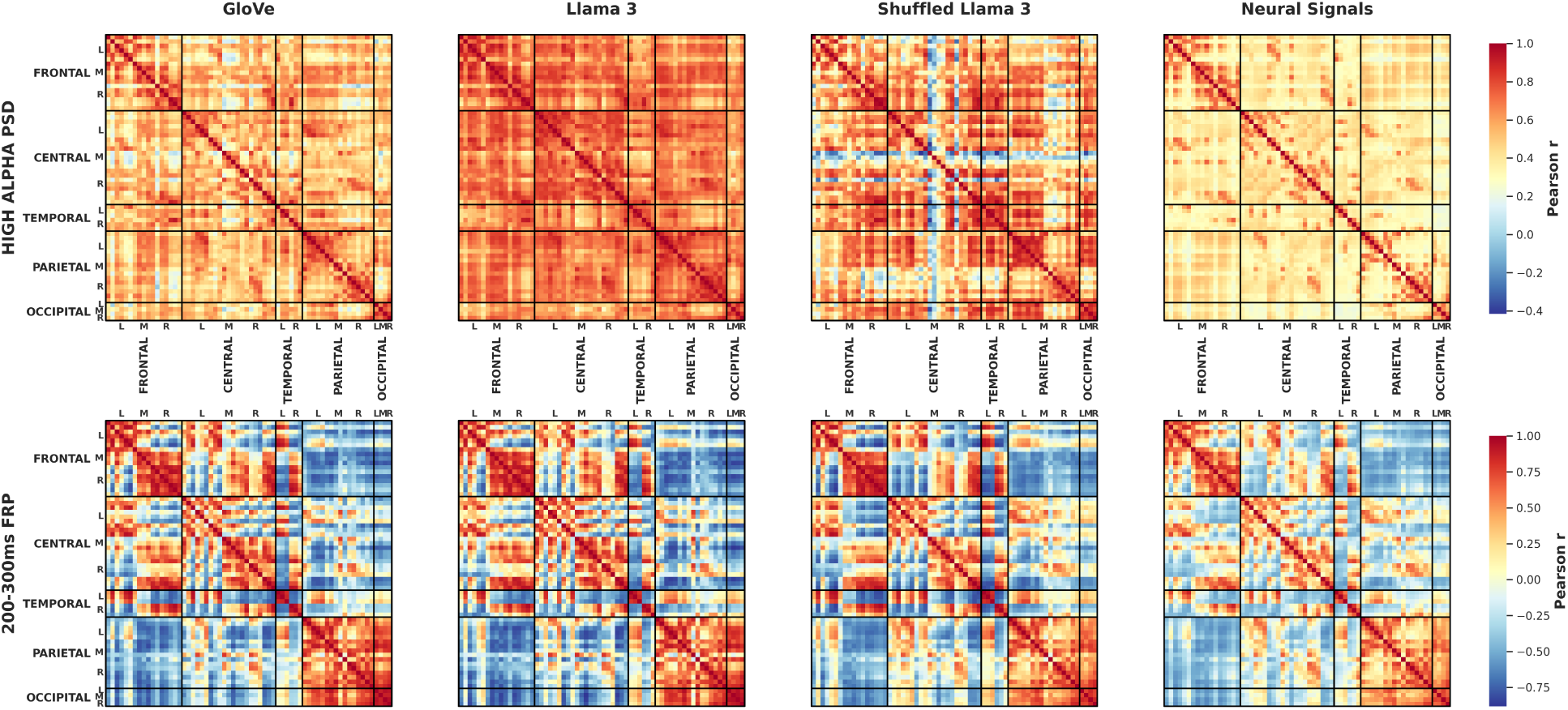
Inter-electrode correlation matrices of recovered encoding-model weights for high-alpha PSD features (top row) and 200–300 ms FRP features (bottom row) derived from GloVe, Llama 3, and shuffled Llama 3 embeddings, alongside the inter-electrode correlation structure of the corresponding raw neural features. Electrodes are grouped by cortical region and hemisphere. Within each row, all matrices share a common color scale representing Pearson correlation coefficients (r).

Across embedding models and neural features, electrodes within the same cortical region generally exhibited stronger correlations than electrodes from different regions. This pattern was observed for both PSD and FRP features. Overall, model weights were more strongly correlated across electrodes for PSD features than for FRP features. For PSD features, Llama 3 exhibited higher overall inter-electrode correlations than GloVe, whereas FRP features showed similar correlation patterns across the two embedding models. We also observed lateralized patterns within the frontal, central, and temporal regions. For PSD features, correlations were generally stronger among electrodes within the same hemisphere than across hemispheres. In contrast, FRP features exhibited positive within-hemisphere correlations and negative correlations between hemispheres.

The inter-electrode correlations derived from the raw neural features exhibited a large-scale spatial organization similar to that observed in the encoding-model weights, including stronger correlations within cortical regions than between regions. For PSD features, the overall spatial patterns were comparable, although the correlation magnitudes differed between the encoding-model weights and the raw neural features. Shuffled Llama 3 embeddings retained many of the broad organizational patterns observed in the original Llama 3 results. For FRP features, the shuffled and original Llama 3 weights showed highly similar inter-electrode correlation structures, both of which closely matched the organization observed in the raw neural features. In contrast, for PSD features, the shuffled Llama 3 weights displayed several correlation patterns that differed from those observed in both the original Llama 3 weights and the raw neural features.

### Semantic Interpretation of Encoding Model Weights

To examine the semantic information captured by the encoding models, we projected the recovered weights back into the embedding space and identified the highest- and lowest-scoring words associated with each neural feature within each brain region. For visualization purposes, we present the Llama 3 results from the parietal region, which exhibited the strongest encoding performance, for high-alpha PSD and 200–300 ms FRP features in Table 1.

**Table 1.** Highest- and lowest-scoring words obtained by projecting recovered Llama 3 encoding-model weights back into the embedding space. Results are shown for the parietal high-alpha PSD feature and the central 200–300 ms FRP feature.

| Neural Feature | Brain Region | Score | Top 20 Words |
| --- | --- | --- | --- |
| High-alpha PSD | Parietal | Highest | words, text, many, copies, repetitions, appears, occur, transcription, populations, Words, text, repeat, indications, divisions, various, yield, scores, contains, illustrations, strategy |
|  |  | Lowest | Williams, 26, Association, player, 8, born, male, Pluto, female, AU, 27%, September, Price, minor-planet, time, players, eccentric, occasions, Graf, world |
| 200–300 ms FRP | Central | Highest | islands, 1940s, two, rosettes, 9.3, 1930s, 1897, 1940s, causeways, roots, 1932, approximately, 1930s, 1870, 1896, 1895, 1920s, 1962, 1898, 1914 |
|  |  | Lowest | although, attributed, involved, seen, given, even, made, although, serve, directed, comes, used, passed, brought, known, consisting, another, claims, worked, showing |

MW, several electrodes exhibited weak or even negative correlations between predicted and observed neural responses (Figure 5). Consequently, subtracting MW performance from normal-reading performance revealed widespread clusters of electrodes showing significantly stronger brain–language alignment during attentive reading.

For the high-alpha PSD feature, positively weighted words included terms related to strategic actions, interpretation, and reporting, whereas negatively weighted words included object names, quantities, and proper nouns. For the 200–300 ms FRP feature, positively weighted words primarily consisted of dates, numbers, and historical references, whereas negatively weighted words included action verbs and other semantically related lexical items.

### Effects of Mind-Wandering on Brain–Language Alignment

We next investigated whether mind wandering altered brain–language alignment.

Encoding models were trained using balanced datasets containing equal numbers of normal-reading and MW fixations and were subsequently evaluated separately on held-out normal-reading and MW samples.

Figure 5 shows that every embedding model exhibited an increase in the number of significantly predicted neural targets when evaluated on normal-reading fixations. For PSD, GPT-2 demonstrated the largest difference between conditions, with 135 channel-by-frequency targets showing significantly stronger encoding performance (i.e., higher correlations between predicted and observed neural responses) during normal reading than during mind-wandering.

**Figure 5.**
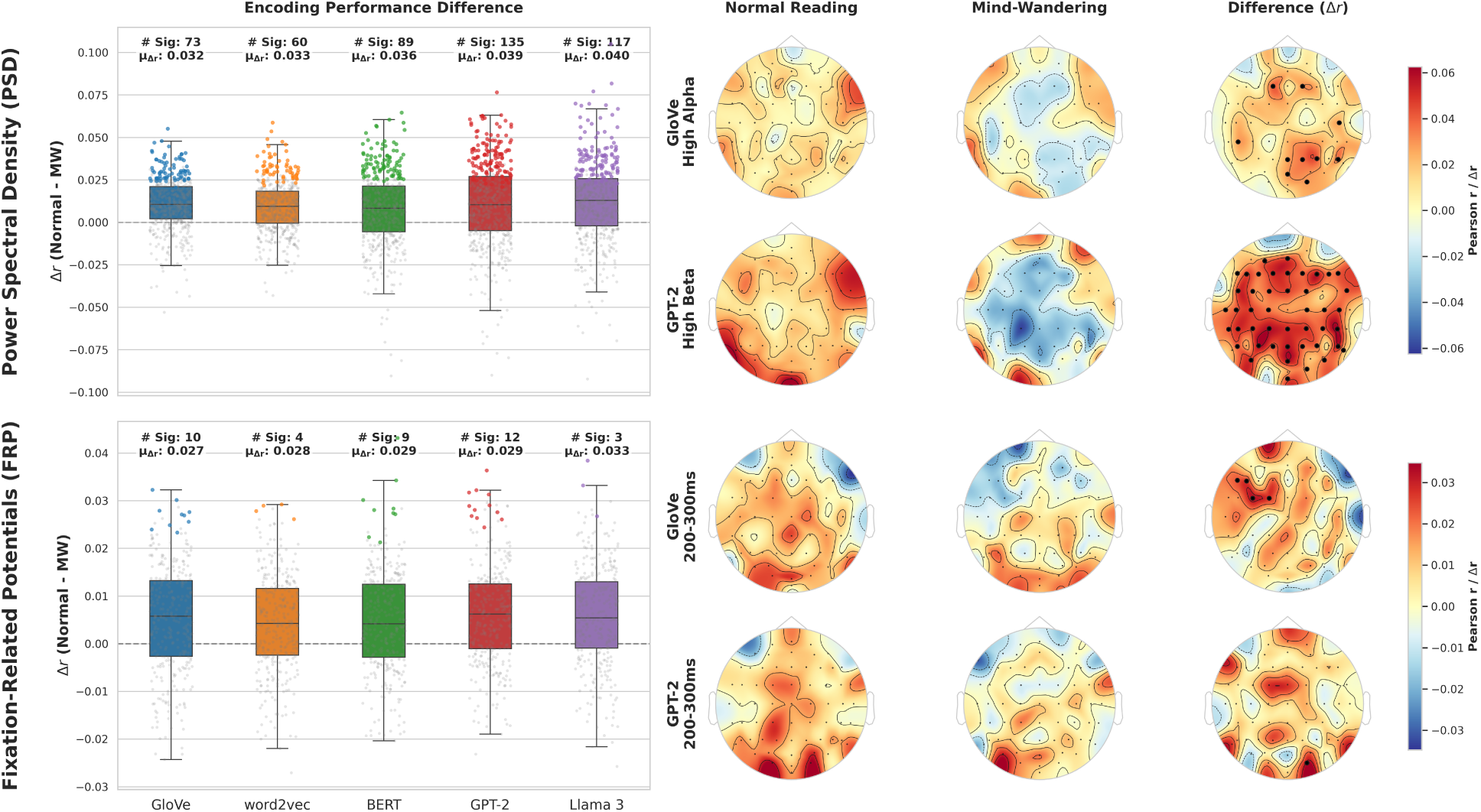
Effects of mind wandering on brain–language alignment. Left panels show the distribution of differences in encoding performance for significant neural features across embedding models. Numbers above boxplots indicate the number of neural features exhibiting significantly stronger encoding performance during normal reading than during MW (p < 0.01, FDR corrected), along with the mean performance difference. Positive values indicate stronger brain–language alignment during normal reading. Right panels show topographic maps of encoding performance during normal reading and mind wandering, as well as their difference. Color indicates the Pearson correlation, and black dots denote electrodes exhibiting significant differences between conditions.

Topographic analyses revealed that the largest MW-related reductions occurred within the high-alpha and low-beta frequency bands (Supplementary Materials). During MW, several electrodes exhibited weak or even negative correlations between predicted and observed neural responses (Figure 5). Consequently, subtracting MW performance from normal-reading performance revealed widespread clusters of electrodes showing significantly stronger brain–language alignment during attentive reading.

In contrast, the effects of MW on FRP-based encoding were substantially weaker. Although encoding performance during MW was generally reduced, the overall spatial distribution of significant correlations remained similar to that observed during normal reading. For example, the 200–300 ms window continued to exhibit central, parietal, and occipital encoding effects during MW. As a result, relatively few electrodes showed statistically significant differences between attentional states.

## Discussion

### Main Findings

In this study, we investigated the relationship between linguistic representations and neural activity during naturalistic reading using an encoding-model framework. By predicting fixation-aligned EEG features from both static and contextual word embeddings, we examined whether information contained within computational language representations is reflected in ongoing neural activity. Consistent with our first hypothesis, significant brain–language alignment was observed across both spectral and event-related EEG measures. Furthermore, consistent with our second hypothesis, this alignment was systematically reduced during mind wandering, suggesting that attentional disengagement weakens the coupling between linguistic input and neural processing. Together, these findings provide evidence that language-model representations capture information reflected in human neural activity during reading and demonstrate that attentional state plays an important role in shaping this relationship.

We employed permutation testing to assess statistical significance and applied false discovery rate correction to account for multiple comparisons, ensuring that the observed effects were unlikely to arise from chance alone. Under these stringent statistical criteria, we found that linguistic representations derived from modern language models significantly predicted fixation-aligned EEG responses during naturalistic reading. The observed correlation coefficients were relatively modest in magnitude compared to those reported in ECoG studies (Goldstein et al., 2022, 2024, 2025a), but this reduction was expected given the complexity of the task and the inherently noisy nature of EEG recordings. Unlike controlled laboratory paradigms that isolate specific linguistic manipulations, naturalistic reading involves substantial variability arising from fluctuations in attention, comprehension, eye movements, and other ongoing cognitive processes (Rayner, 1998).

Consequently, even modest but reliable correlations provide meaningful evidence that linguistic information is represented in low signal-to-noise ratio neural signals such as EEG.

We found that, for both PSD and FRP features, contextual embeddings generally achieved stronger encoding performance than static embeddings, as evidenced by a greater number of significantly predicted neural features and higher average correlation coefficients. Consistent with previous studies (Caucheteux & King, 2022; Goldstein et al., 2022, 2024), this superiority of contextual embeddings suggests that neural activity during reading reflects more than isolated lexical properties and is sensitive to higher-order contextual information captured by modern language models. Among the contextual embeddings examined, Llama 3 generally produced the strongest encoding performance. This may be due to its substantially larger model size relative to GPT-2 and BERT, which provides greater representational capacity for capturing complex semantic relationships in language (Antonello et al., 2023). However, the present study was not designed to directly evaluate the contribution of model size or architectural differences, and future work is needed to determine the extent to which these factors account for differences in brain–language alignment. Importantly, these findings indicate that higher-order semantic representations, which are better captured by contextual embeddings, remain detectable even in low-spatial-resolution neural recordings such as EEG during naturalistic reading.

For PSD features, we observed the strongest brain–language alignment in the alpha and low-beta frequency ranges. Alpha oscillations have long been associated with attentional allocation and semantic processing during language comprehension (Kelly et al., 2006; Klimesch, 2012; Kornrumpf et al., 2017; Ray & Cole, 1985; Röhm et al., 2001). During reading, words vary in their semantic complexity, predictability, and contextual integration demands, which in turn influence the attentional resources required for processing. Therefore, word embeddings, which capture many of these linguistic properties, may share variance with alpha-band activity. In addition, recent work has shown that alpha oscillations coordinate information exchange between the oculomotor and visual systems during natural reading, and this coordination becomes more pronounced for demanding words (Pan et al., 2023). Because eye-movement behavior, including fixation durations and saccade patterns, is also influenced by linguistic properties (Kliegl et al., 2004; Pan et al., 2023), this mechanism may further contribute to the strong brain–language alignment observed in the alpha band. Previous studies have linked beta oscillations to several aspects of language processing (Weiss & Mueller, 2012). In particular, beta activity has been proposed to support top-down predictions of upcoming stimuli during language comprehension (Arnal et al., 2011; Bressler & Richter, 2015; Fujioka et al., 2009; Richter et al., 2017) and has been implicated in semantic integration and memory-related processes (Bastiaansen et al., 2010; Weiss & Rappelsberger, 2000; Weiss & Mueller, 2003). Because word embeddings encode semantic and contextual information, they may share variance with these beta-related language functions. This relationship may be particularly strong for contextual embeddings, which capture predictive and contextual information beyond individual word meanings, potentially explaining their superior encoding performance in the beta band.

In contrast, we observed relatively little brain–language alignment within the gamma frequency range. This finding is consistent with a previous study of naturalistic reading using MEG, which reported alpha- and beta-band effects but comparatively limited gamma-band effects (Mäkelä et al., 2024). One possible explanation is that language-related gamma effects may occur at higher frequencies than those examined in the present study. For example, successful semantic encoding has been associated with increases in gamma power within the 55–70 Hz range (Hanslmayr et al., 2008), whereas our analyses were restricted to frequencies below 50 Hz. Another possible explanation is the lower signal-to-noise ratio of scalp-recorded gamma activity, which is particularly susceptible to muscular and ocular artifacts (Hipp & Siegel, 2013). As a result, language-related gamma activity may be more difficult to detect using scalp EEG than lower-frequency oscillations.

For the 1/f-corrected PSD analysis, the uncorrected PSD features often exhibited numerically higher brain–language alignment but rarely reached statistical significance over peripheral electrodes. In free-viewing EEG recordings, the aperiodic 1/f component may reflect a combination of neural and non-neural contributions, including broadband activity arising from muscle and ocular artifacts that are often strongest at peripheral electrodes (Gerster et al., 2022). As a result, 1/f correction reduces broadband spectral noise and enhances the specificity of oscillatory activity estimates. Nevertheless, the spatial distribution and number of significant encoding effects were largely comparable between the corrected and uncorrected analyses, which reflects the robustness of the permutation testing and FDR-correction procedures. Given the similarity of the overall results, we elected to use the uncorrected PSD features for subsequent analyses, including the comparison between normal reading and mind wandering. One consideration is that the 1/f baseline, although estimated from the entire reading session, may still contain characteristics associated with mind wandering, particularly for participants who reported mind wandering on many reading pages. Consequently, the correction procedure could remove variance that is directly relevant to the attentional-state effects examined in the present study.

We observed strong brain–language alignment in FRP features beginning approximately 100 ms after fixation onset, with the strongest effects occurring between 200 and 300 ms. Although the P1 component exhibited the largest amplitude in the grand-average FRPs, relatively weak encoding performance was observed during the 0–100 ms interval. This finding is not surprising, as the P1 component is primarily associated with early visual processing and has been shown to be modulated by parafoveal preview and visual word form (Henderson et al., 2013; Yagi, 1981). Since these processes are indirectly related to semantic information represented by word embeddings, relatively weak brain–language alignment would be expected for this early stage of FRPs. Encoding performance increased during the 100–200 ms interval, corresponding approximately to the N1 component. Previous studies have linked activity within this time window to orthographic and phonological processing, including the activation of lexical representations and sensitivity to word frequency and parafoveal preview. Since these processes are more directly related to lexical properties than the earlier visual components, stronger correspondence between word embeddings and FRPs may emerge during this period.

The strongest encoding effects were observed between 200 and 300 ms, particularly over central and parietal-occipital electrodes. This interval overlaps with the P2, N2, and P300 components (Degno & Liversedge, 2020). Specifically for the P2 component, previous studies have reported modulation by semantic relatedness and word predictability (Baccino & Manunta, 2005; Kretzschmar et al., 2015). Although relatively few studies have examined the P2 component and its functional role remains not fully understood (Degno & Liversedge, 2020), our results suggest that neural activity within this interval contains substantial information related to the linguistic representations, and this brain–language alignment emerges before the classical N400 period during naturalistic reading. This result is consistent with findings from a previous study using Japanese narratives, in which semantic vectors derived from the relationship between the original text and participants’ content reports were shown to predict N1, P2, and P3 responses (Sato & Mizuhara, 2018).

Interestingly, encoding performance remained significant during the 300–500 ms interval (i.e., the N400 component) but was generally weaker than that observed between 200 and 300 ms. The N400 is one of the most extensively studied language-related ERP components (Degno & Liversedge, 2020) and has been shown to be sensitive to a wide range of linguistic factors, including parafoveal preview, semantic relatedness, word predictability, and syntactic or semantic violations (Degno et al., 2019; Dimigen et al., 2012; Kornrumpf et al., 2016; López-Peréz et al., 2016; Metzner et al., 2017). Given its well-established role in language comprehension, we would expect strong brain–language alignment during this time window, consistent with previous encoding studies that reported robust relationships between language representations and N400 activity (Sassenhagen & Fiebach, 2020; Schwartz & Mitchell, 2019). However, the encoding effects observed during the N400 period were weaker than those associated with earlier FRP components, which may be due to the difference in experimental paradigm. Prior studies used highly controlled word-by-word or sentence-level reading tasks (Sassenhagen & Fiebach, 2020; Schwartz & Mitchell, 2019), whereas we examined naturalistic paragraph reading. Under natural reading conditions, later FRP components are more likely to overlap with responses elicited by subsequent fixations, potentially reducing the separability of N400-related activity. Future work could address this limitation using deconvolution approaches (Ehinger & Dimigen, 2019) to better isolate overlapping FRPs.

Overall, FRP-based encoding produced stronger correlations and a greater number of significant neural features than PSD-based encoding. We interpret this finding as evidence that FRPs reflect neural responses that are directly time-locked to visual word input and the subsequent stages of lexical and semantic processing (Degno & Liversedge, 2020). In contrast, PSD features reflect ongoing neural dynamics that may be influenced by broader cognitive factors, including attention, contextual integration, and linguistic prediction (Bastiaansen & Hagoort, 2006). As a result, the relationship between individual words and neural activity may be more direct for FRPs than for PSD features. This difference may also explain why the performance gap between static and contextual embeddings was smaller for FRP features than for PSD features. Since both static and contextual embeddings contain substantial information about individual words, strong encoding performance can be achieved even without rich contextual representations. In contrast, contextual embeddings may provide a greater advantage for PSD features by capturing higher-order contextual information that contributes to ongoing neural activity during reading (Hollenstein et al., 2019).

Because individual model coefficients are difficult to interpret directly (Kriegeskorte & Douglas, 2019), we examined the similarity structure of the learned model weights across electrodes. Model weights are influenced by both the input representations (word embeddings) and the output neural signals, so similarities in weight patterns can provide insight into the factors driving brain–language alignment (Haufe et al., 2014). We found that the choice of word embedding model had relatively little impact on the overall weight-similarity structure. In contrast, substantial differences were observed between neural feature types. Specifically, model weights were highly correlated across electrodes for PSD features but exhibited more localized patterns for FRP features. This finding is consistent with the nature of the neural signals themselves: PSD features reflect ongoing oscillatory activity that is often distributed across large-scale neural networks (Buzsaki & Draguhn, 2004), whereas FRPs capture transient responses time-locked to individual fixation events and are therefore more spatially specific (Degno & Liversedge, 2020; Dimigen et al., 2011). Together, these results suggest that the structure of the learned model weights is influenced more strongly by the properties of the neural signals than by differences among word embedding models. This interpretation is reasonable given that all embedding models encode broadly similar semantic information, whereas neural activity varies substantially across electrodes, frequency bands, and temporal components.

Nevertheless, contextual embeddings produced more globally correlated weight patterns than static embeddings in the alpha and low-beta bands, suggesting that contextual information contributes additional explanatory power beyond lexical semantics alone.

To gain further insight into the semantic dimensions captured by the encoding models, we projected the recovered model weights back into the embedding space and identified the highest- and lowest-scoring words associated with brain regions containing a large number of electrodes that exhibited significant encoding effects for a given neural feature. For parietal high-alpha PSD features, positively weighted words included terms related to strategic actions, interpretation, and reporting, which may require greater contextual integration and semantic interpretation. In contrast, negatively weighted words consisted primarily of object names, quantities, and proper nouns that can often be understood with relatively limited contextual information. Given the established role of alpha oscillations in attention and semantic processing (Kelly et al., 2006; Klimesch, 2012; Kornrumpf et al., 2017; Ray & Cole, 1985; Röhm et al., 2001), these differences may contribute to the stronger association between such semantic dimensions and ongoing oscillatory activity. Interestingly, for FRP features within the 200–300 ms interval, positively weighted words consisted primarily of numbers. One possible explanation is that numbers are relatively infrequent compared to ordinary words and may therefore elicit enhanced attentional responses, potentially resembling oddball-like effects associated with the pre-P300/P300 period (Sutton et al., 1965). Alternatively, numerical information may engage partially distinct cognitive processes compared to ordinary lexical-semantic processing (de Chambrier et al., 2023). However, the present analysis was intended primarily as an exploratory examination of the semantic dimensions captured by the encoding models rather than a definitive characterization of their linguistic content. Future work should investigate these semantic patterns more systematically using quantitative linguistic measures such as word frequency, surprisal, and contextual complexity to provide a clearer understanding of the linguistic factors that contribute most strongly to brain–language alignment during naturalistic reading.

One major novelty of the ROAMM dataset (Sun et al., 2026b), beyond its naturalistic reading paradigm, is the availability of span-level MW annotations throughout the entire reading period. This allowed us to investigate a question that has been largely overlooked in previous brain–language studies: how does attentional state influence the relationship between linguistic input and neural activity? To address this question, we constructed balanced datasets containing equal numbers of normal-reading and MW fixations to eliminate sample size as a potential confound. Encoding models were then trained on this balanced dataset and evaluated separately on normal-reading and MW samples. Across embedding models, encoding performance was consistently stronger during normal reading than during MW (Chen et al., 2025), supporting our hypothesis that attentional disengagement weakens brain–language alignment.

Interestingly, we found that the effects of MW were considerably stronger for PSD features than for FRP features. For PSD, significant reductions in encoding performance were particularly evident within the alpha and beta frequency bands. Previous studies have shown that both frequency ranges are strongly modulated by MW (Braboszcz & Delorme, 2011; Jin et al., 2019; Kam et al., 2022). According to decoupling theory (Smallwood & Schooler, 2006), MW redirects cognitive resources away from external stimuli and toward internally generated thoughts. During attentive reading, neural activity remains closely coupled to the processing of incoming linguistic information (Klimesch, 2012; Weiss & Mueller, 2012). In contrast, during MW, a larger proportion of neural activity may reflect internally generated cognitive processes unrelated to the text. As a result, activity in these bands may simultaneously reflect both language-related processing and fluctuations in attentional state, reducing the amount of variance that can be explained by word embeddings and consequently weakening brain–language alignment.

In contrast, FRP-based encoding performance remained relatively stable across attentional states. Although previous studies have reported reductions in FRP components such as the P1, N1, and P300 during MW (Dong et al., 2021; Jin et al., 2019; Kam et al., 2022), these effects were not significant in the context of the present encoding analysis. We believe there are at least two possible explanations for this finding. First, MW during naturalistic reading may primarily disrupt higher-order cognitive processes (Smallwood et al., 2008) while leaving lower-level visual and lexical processing relatively intact.

Readers experiencing MW often continue moving their eyes through the text and processing individual words (Sun et al., 2026a), even though comprehension and sustained attention are reduced. In this sense, MW during reading may be characterized more by a loss of the overall narrative thread than by a complete failure to process individual words. Second, certain linguistic properties of the text may themselves contribute to the onset or maintenance of MW. Difficult, repetitive, or less engaging content may increase the likelihood of attentional disengagement (Feng et al., 2013). As a result, changes in FRPs associated with MW may occur in the same direction as changes associated with the linguistic properties of the words themselves, leading to a relatively small reduction in encoding performance. We further argue that this same-direction relationship is less likely to occur for PSD features. Unlike the relatively consistent attenuation observed in FRPs, PSD features during MW may reflect a mixture of linguistic, attentional, and internally generated cognitive processes that do not necessarily vary in the same direction.

Taken together, these findings suggest that MW primarily affects brain–language alignment through changes in ongoing neural states rather than through a complete disruption of word-level processing. Ongoing oscillatory activity appears particularly sensitive to fluctuations in attention, whereas fixation-related potentials remain more closely tied to the processing of individual words. More broadly, these results indicate that neural features that are directly linked to stimulus events may provide more robust measures of stimulus–brain relationships under naturalistic conditions. For studies examining the relationship between language representations and neural oscillations, these findings demonstrate that periods of attentional disengagement can substantially weaken measured brain–language alignment, suggesting that attention may represent a largely overlooked source of variability. Future encoding studies may therefore benefit from incorporating measures of attentional state when investigating the neural representation of language.

### Limitations and Future Directions

Several limitations should be considered when interpreting the present findings. While we demonstrated significant relationships between word embeddings and regional neural signals, the findings were derived from an encoding-model framework rather than direct analyses of neural representations. Consequently, the observed effects should be interpreted as evidence that linguistic representations can predict variance in neural activity. The identified frequency bands and time windows that were most predictable from word embeddings do not necessarily reflect the neural mechanisms that generate language representations. Future work employing raw and source-localized neural representations may provide a more direct characterization of these mechanisms.

The use of the ROAMM dataset (Sun et al., 2026b) and its naturalistic free-viewing reading paradigm introduces additional considerations. While this design provides high ecological validity and allows investigation of language encoding under conditions that more closely resemble everyday reading, it also introduces challenges that are absent in tightly controlled word-by-word paradigms. Readers were free to make regressions and multiple fixations on the same word. In the present study, we treated each fixation as an independent sample regardless of whether the word had been previously fixated. Because regressions may reflect rereading, memory retrieval, or other cognitive states (Madureira et al., 2023; Schotter et al., 2014), neural responses during regression fixations may differ from those observed during first-pass reading. Future work should explicitly separate first-pass and regression fixations to determine how rereading influences brain–language alignment.

Parafoveal and peripheral processing present another source of uncertainty. Previous studies have demonstrated that parafoveal information affects fixation-related neural responses during reading (Dimigen et al., 2012; Henderson et al., 2013; Yagi, 1981).

In the present study, we derived linguistic features from the currently fixated word. Although contextual embeddings partially incorporate information from surrounding text through their larger context windows, the extent to which contextual embeddings account for parafoveal influences remains unclear.

We also acknowledge several limitations related to the feature-extraction procedures used in the present study. Because little prior work has examined brain–language alignment using fixation-level EEG spectral and fixation-related potential features, we adopted relatively general approaches for defining temporal windows. For PSD analyses, spectral power was extracted from fixation-limited windows. However, language-related neural processes may emerge only after a delay following fixation onset, creating potential temporal misalignment between the extracted features and the underlying cognitive processes. Although semantic processing and oscillatory power may evolve slowly enough to preserve substantial overlap despite this misalignment, the precise temporal relationship remains unclear. For FRP analyses, we summarized neural activity using non-overlapping 100 ms windows. While this approach was motivated by the temporal characteristics of established FRP components, it may not fully capture the precise timing of language-related neural responses. Future work could employ sliding-window approaches or higher-temporal-resolution analyses to more precisely characterize the temporal dynamics of brain–language alignment.

Consistent with previous studies, we extracted contextual embeddings from the final hidden layers of the language models (Goldstein et al., 2024; Hollenstein et al., 2019), as prior work using such models has typically found that mid-to-late layers best predict brain activity (Caucheteux & King, 2022). However, prior work using ECoG recordings has reported a relationship between language-model depth and the brain’s temporal receptive window, with middle layers exhibiting stronger brain–language alignment than either earlier or later layers (Goldstein et al., 2025a). Whether similar layer-specific effects exist for EEG remains unknown. Furthermore, the optimal layer may vary across model architectures and neural features. Future studies should systematically evaluate representations from different layers to determine which levels of language-model processing best correspond to EEG activity during reading.

We further reduced the dimensionality of the embedding representations using PCA prior to model fitting. This approach substantially reduced computational requirements and enabled comparisons across multiple embedding models. However, components explaining relatively little variance may still contain information relevant to neural activity. Dimensionality reduction may therefore remove information that contributes to brain–language alignment. It may also reduce differences between language models with substantially different embedding dimensionalities. For example, reducing all embedding spaces to 50 principal components may partially diminish advantages associated with larger embedding spaces such as those of Llama 3. Future work could investigate encoding performance using the original embedding spaces or employ alternative dimensionality-reduction methods that better preserve information relevant to neural responses.

The encoding framework optimized model parameters using mean squared error (MSE) while evaluating performance using Pearson correlation. This approach is common in neural encoding studies and facilitates direct comparison with previous work (Chen et al., 2025; Goldstein et al., 2022, 2024, 2025a; Hosseini et al., 2024). However, MSE and correlation quantify different aspects of prediction quality. A model may achieve high correlation with the target signal while exhibiting relatively large prediction errors in magnitude, or vice versa. As a result, optimizing MSE may not necessarily maximize the evaluation metric used to assess encoding performance. Recent studies have explored various relationship-alignment techniques such as Canonical Correlation Analysis and Wasserstein distance, which directly optimize representational similarity between neural and linguistic features, to learn feature transformations (Qiu et al., 2023). Future work could adapt these approaches and investigate whether correlation-based objectives improve encoding performance and provide a more direct alignment between training and evaluation criteria.

We performed the encoding analyses across participants, which provided substantially more training data and produced models that were less susceptible to noise and outliers. This approach also allowed the resulting encoding models to capture patterns that are more representative of the general population. However, it assumes that the relationship between linguistic representations and neural responses is relatively consistent across individuals. While reading engages broadly similar visual and language-processing systems across readers (Rayner et al., 2012), previous fMRI studies have reported meaningful individual differences in neural responses during reading (Jangraw et al., 2023).

Future work should investigate how much data are required to obtain stable participant-specific encoding models and whether individual models reveal patterns that are obscured in group-level analyses.

We treated mind wandering as a binary attentional state because the ROAMM dataset does not provide information regarding the depth, intentionality, or content of individual mind-wandering episodes. Although this enabled a direct comparison between normal reading and mind wandering, these factors may influence neural activity and contribute additional variability within the mind-wandering condition. Future studies incorporating richer experience-sampling measures may provide a more nuanced understanding of how different forms of mind wandering influence the neural representation of language.

Finally, the current study focused exclusively on EEG measures at the fixation level. Because analyses were performed on fixation/word-level samples, many eye-tracking features that unfold across longer temporal intervals, such as saccades, blinks, and higher-order reading behaviors, were not incorporated into the encoding framework. Future work could develop multimodal encoding models that jointly predict neural and oculomotor responses from linguistic representations over longer temporal contexts, such as sentences or paragraphs. Such approaches may provide a more comprehensive characterization of how language is represented across multiple physiological systems and over extended timescales during naturalistic reading.

## Conclusion

The present study demonstrated that linguistic representations derived from modern language models are reflected in fixation-aligned EEG activity during naturalistic reading. Using an encoding-model framework, we found significant brain–language alignment across both spectral and fixation-related potential features. We further showed that different neural features capture language processing, with PSD-based encoding exhibiting the strongest effects in the alpha and low-beta frequency bands and FRP-based encoding peaking within the 200–300 ms interval following fixation onset. By leveraging the unique span-level mind-wandering annotations available in the ROAMM dataset, we also demonstrated that mind wandering weakens the relationship between linguistic representations and neural activity during reading, particularly for ongoing oscillatory activity. Together, these findings extend previous brain–language encoding research from controlled laboratory paradigms to naturalistic reading, provide new insights into the neural correlates of language processing under ecologically valid conditions, and highlight attentional state as an important factor shaping the correspondence between linguistic representations and neural activity.

## Data and Code Availability Statement

The EEG and eye-tracking data supporting this study are part of the openly available ROAMM dataset (Sun et al., 2026b) and can be accessed via OpenNeuro (https://openneuro.org/datasets/ds007629). Mind-wandering annotation and modeling code associated with the ROAMM dataset are described in Sun et al. (2026b) and are publicly available. Analysis code for the encoding models described in this manuscript is available at https://github.com/GlassBrainLab/roamm-brain-language-alignment.

## Supplementary Material

Supplementary materials accompanying this preprint, including complete topographic maps across all embedding models and neural feature types, complete channel-by-channel encoding-model weight similarity matrices, complete topographic maps of mind-wandering effects, the electrode layout used for regional analyses, and representative scatter plots of observed versus predicted neural responses, are provided in a separate file accompanying this preprint.

## Acknowledgements

The authors thank the Vermont Advanced Computing Center (VACC) for computational resources. This research was supported by internal funding from the University of Vermont. The authors have no conflicts of interest to disclose.

