## Supplemental Figures for "Brain–Language Alignment During Naturalistic Reading and Its Disruption by Mind-Wandering"

### Supplementary Material for: Brain–Language Alignment During Naturalistic Reading and Its Disruption by Mind-Wandering

#### Author Note

Correspondence concerning this article should be addressed to Haorui Sun, Department of Electrical and Biomedical Engineering, University of Vermont, Votey Hall, 33 Colchester Ave, Burlington, VT 05405, United States.

#### Abstract

This document provides supplementary figures accompanying the preprint "Brain–Language Alignment During Naturalistic Reading and Its Disruption by Mind-Wandering."

### Supplementary Material for: Brain–Language Alignment During Naturalistic Reading and Its Disruption by Mind-Wandering

This appendix provides supplementary figures that complement the results presented in the main text, including complete topographic maps across all embedding models and neural feature types, complete channel-by-channel encoding-model weight similarity matrices, complete topographic maps of mind-wandering effects, the electrode layout used for regional analyses, and representative scatter plots of observed versus predicted neural responses.

#### Brain–Language Alignment Topographic Maps

#### Model Weight Similarity Matrices

This section provides the complete channel-by-channel weight similarity matrices used in the spatial analysis of encoding-model weights. For each embedding model, frequency band (PSD), or temporal window (FRP), pairwise Pearson correlations were computed between electrode-specific ridge-regression coefficient vectors. Similarity matrices were calculated independently within each cross-validation fold and subsequently averaged across folds. These visualizations summarize the extent to which different scalp regions relied on similar embedding dimensions when predicting neural activity and complement the representative examples presented in the main text.

#### Mind-Wandering Effects on Brain–Language Alignment

#### Supplementary Figures

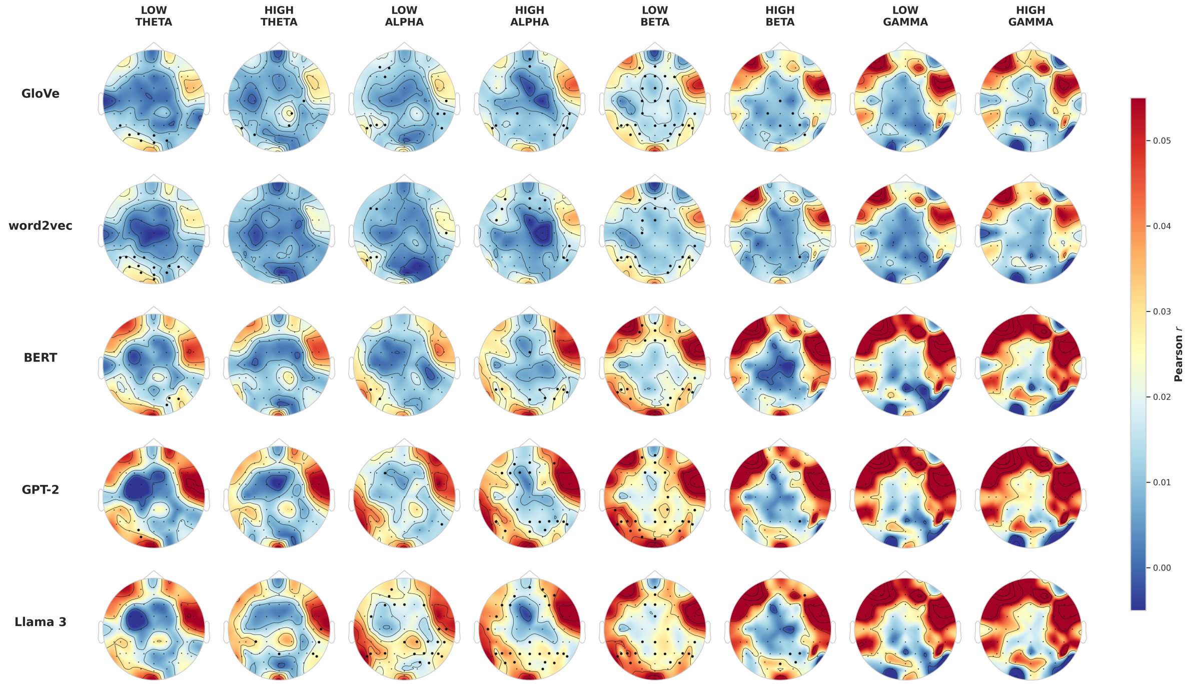


**Figure 1**

*Figure S1. Topographic distributions of encoding performance for fixation-aligned EEG spectral power features across all embedding models and frequency bands. Color indicates the Pearson correlation between predicted and observed neural responses. Black dots denote electrodes exhibiting significant encoding performance ($p < 0.01$, FDR corrected). All panels share a common color scale.*

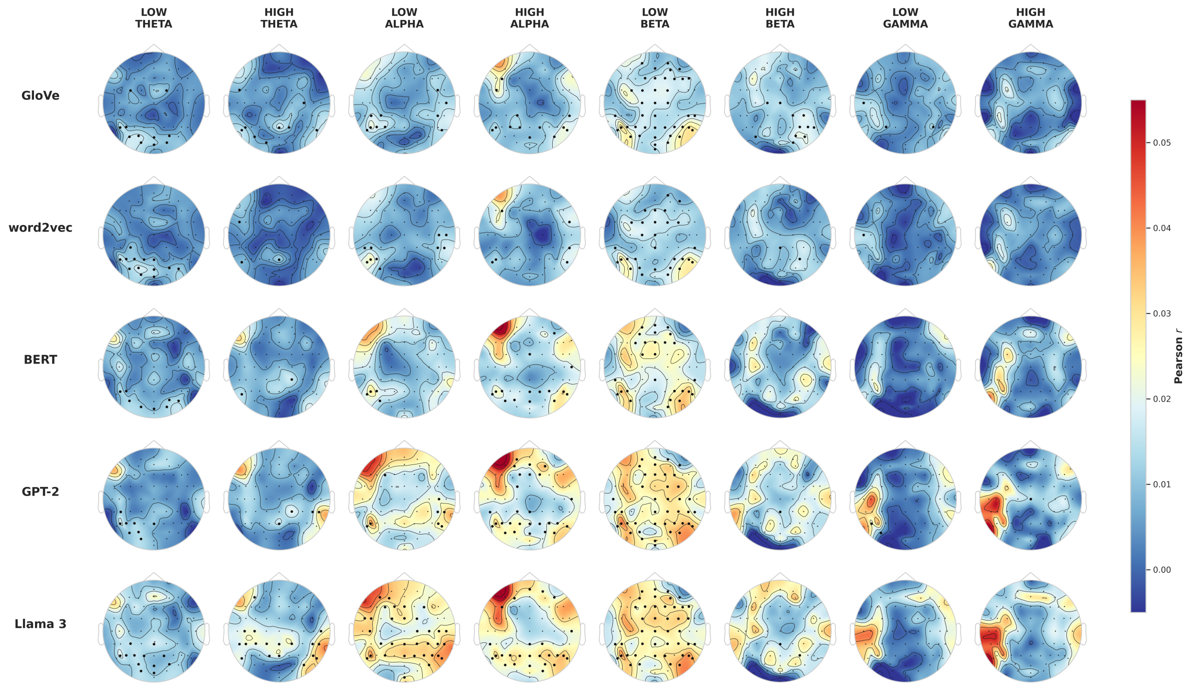


**Figure 2**

*Figure S2. Topographic distributions of encoding performance for fixation-aligned 1/f-corrected EEG spectral power features across all embedding models and frequency bands. Spectral power was normalized relative to channel- and run-specific estimates of the aperiodic spectral background. Color indicates the Pearson correlation between predicted and observed neural responses. Black dots denote electrodes exhibiting significant encoding performance ($p < 0.01$, FDR corrected). All panels share a common color scale.*

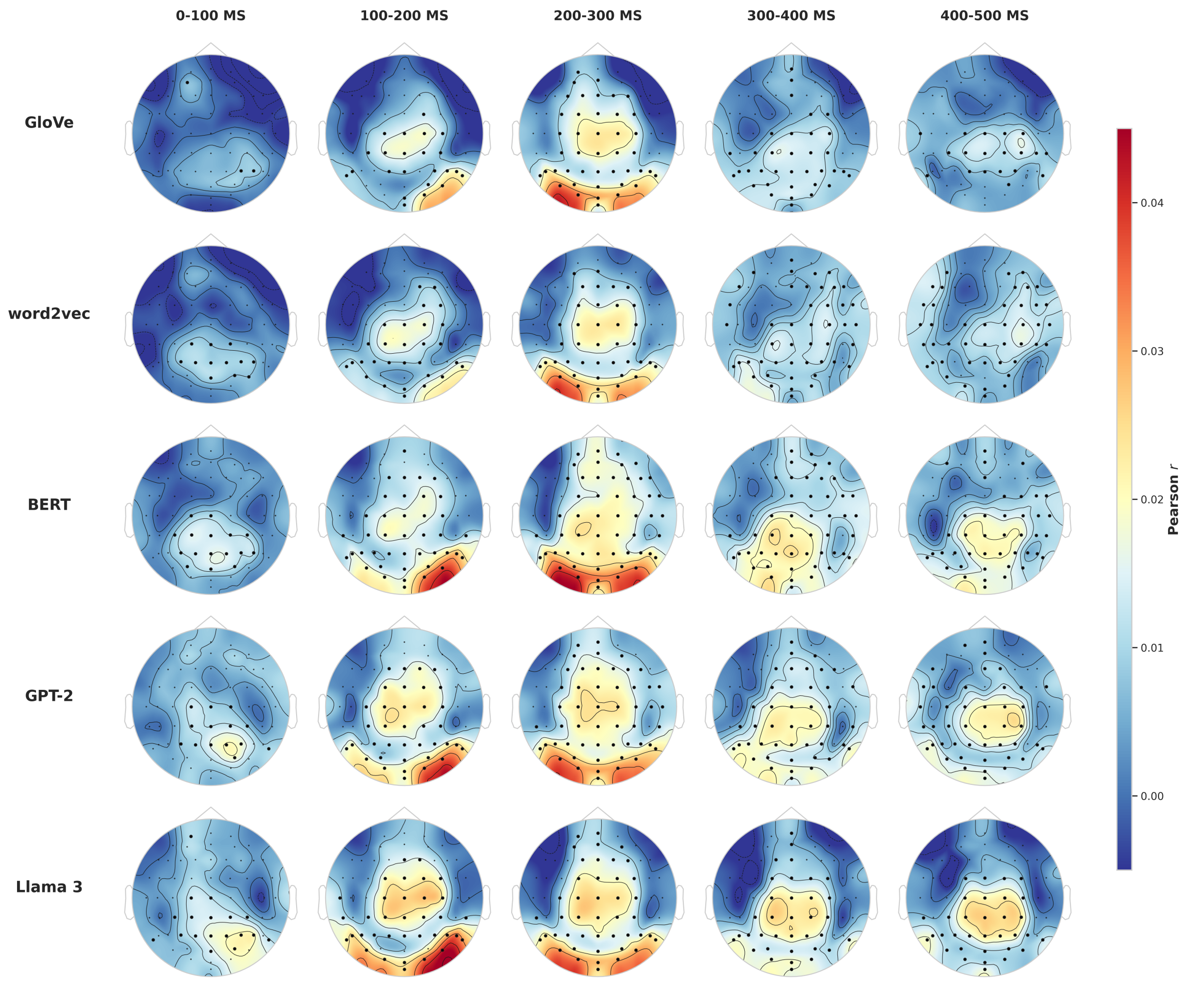


**Figure 3**

*Figure S3. Topographic distributions of encoding performance for fixation-related potential features across all embedding models and post-fixation time windows. Color indicates the Pearson correlation between predicted and observed neural responses. Black dots denote electrodes exhibiting significant encoding performance ($p < 0.01$, FDR corrected). All panels share a common color scale.*

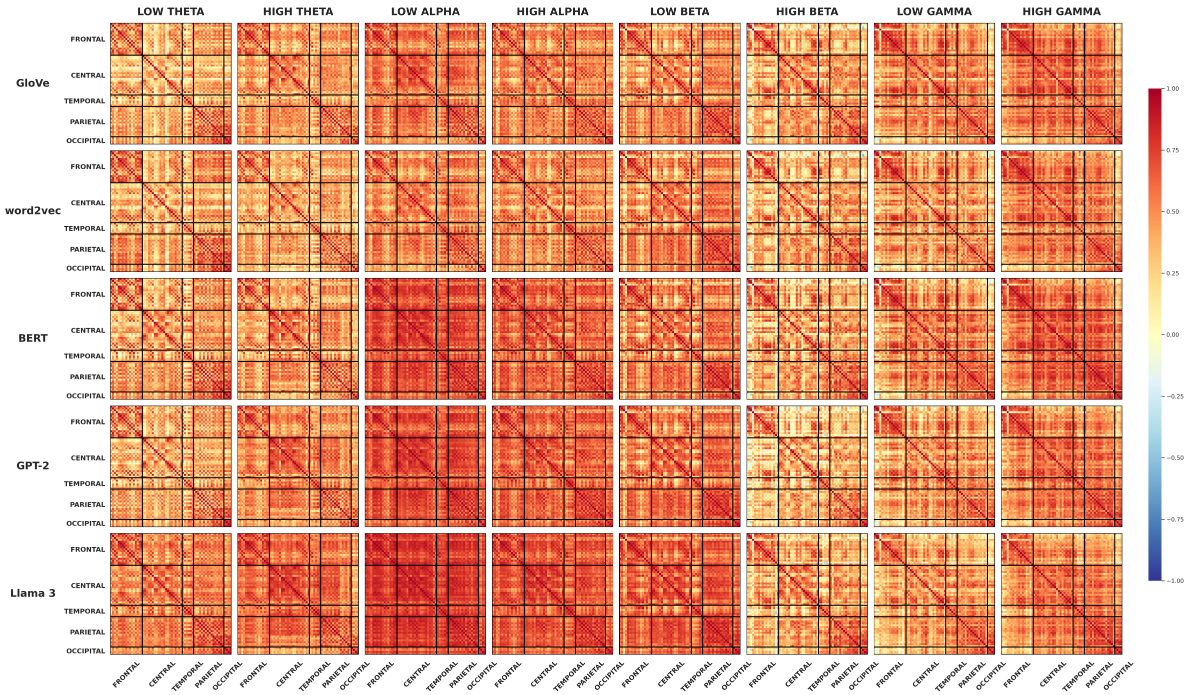


**Figure 4**

*Figure S4. Channel-by-channel similarity matrices for PSD encoding-model weights. Each panel shows the average Pearson correlation between ridge-regression coefficient vectors across electrodes for a specific embedding model and frequency band. Correlations were computed independently within each cross-validation fold and averaged across folds. Electrodes are grouped by scalp region: frontal, central, temporal, parietal, and occipital, with black lines indicating regional boundaries. Warmer colors indicate greater similarity in the embedding dimensions used to predict neural activity.*

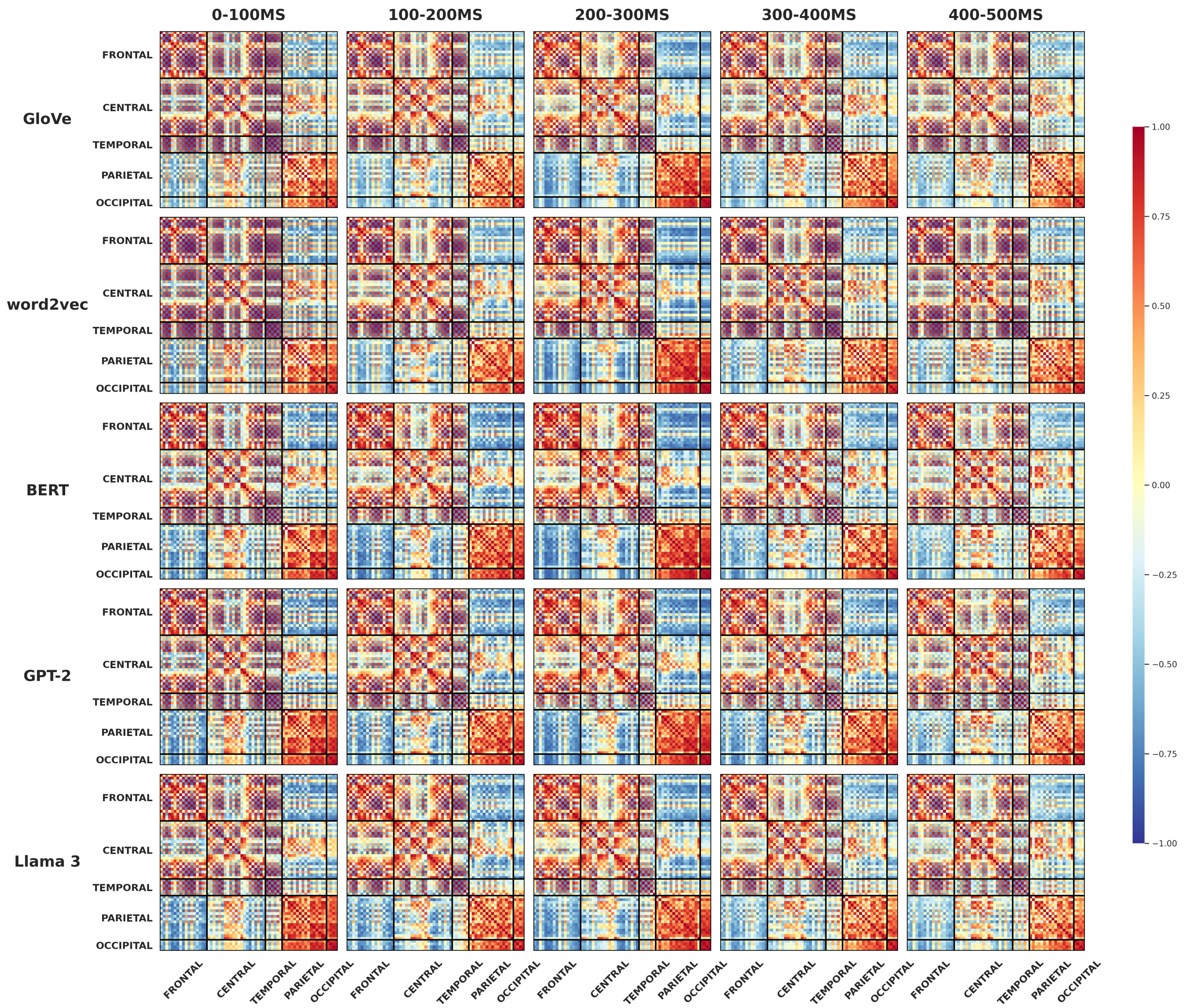


**Figure 5**

*Figure S5. Channel-by-channel similarity matrices for FRP encoding-model weights.*

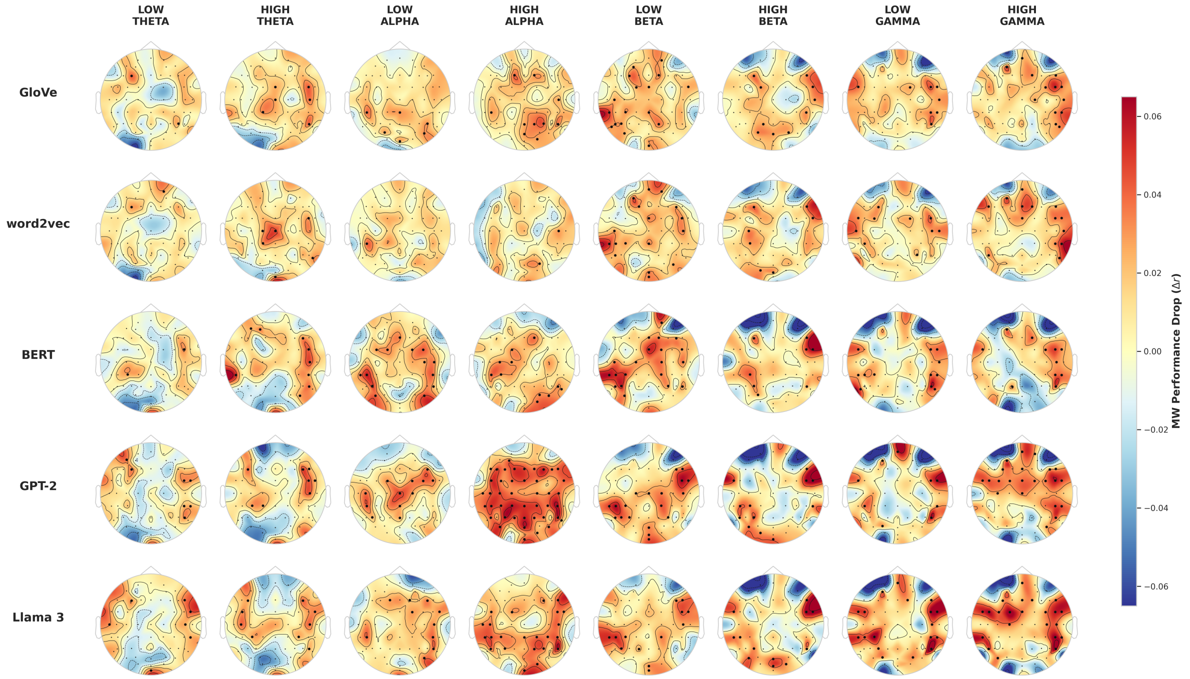


**Figure 6**

*Figure S6. Topographic distributions of differences in encoding performance between normal reading and MW for fixation-aligned EEG spectral power features across all embedding models and frequency bands. Color indicates the difference in Pearson correlation between predicted and observed neural responses. Positive values indicate stronger brain–language alignment during normal reading and vice versa. Black dots denote electrodes exhibiting significant differences between conditions ($p < 0.01$, FDR corrected). All panels share a common color scale.*

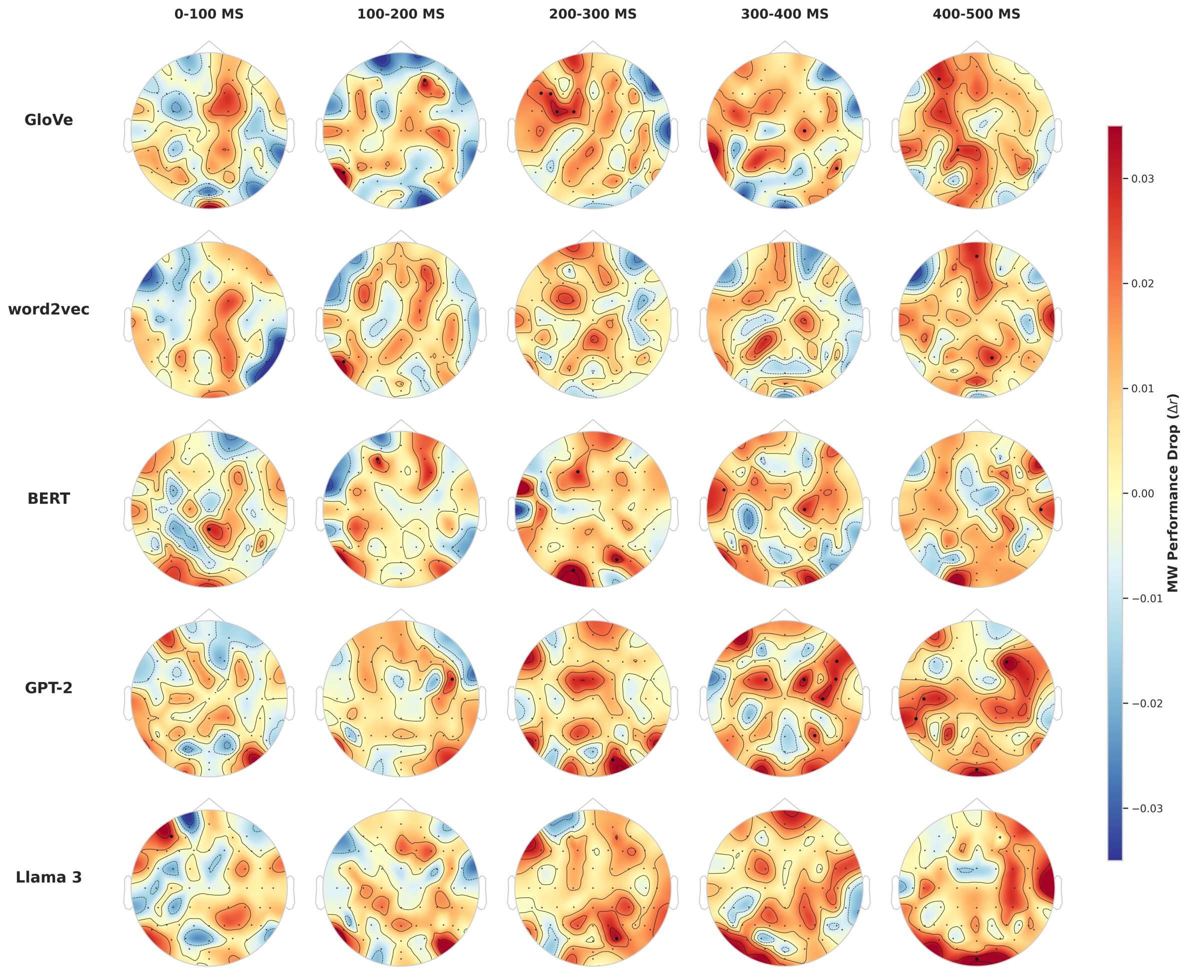


**Figure 7**

*Figure S7. Topographic distributions of differences in encoding performance between normal reading and MW for fixation-related potential features across all embedding models and post-fixation time windows. Color indicates the difference in Pearson correlation between predicted and observed neural responses. Positive values indicate stronger brain–language alignment during normal reading and vice versa for negative values. Black dots denote electrodes exhibiting significant differences between conditions ($p < 0.01$, FDR corrected). All panels share a common color scale.*

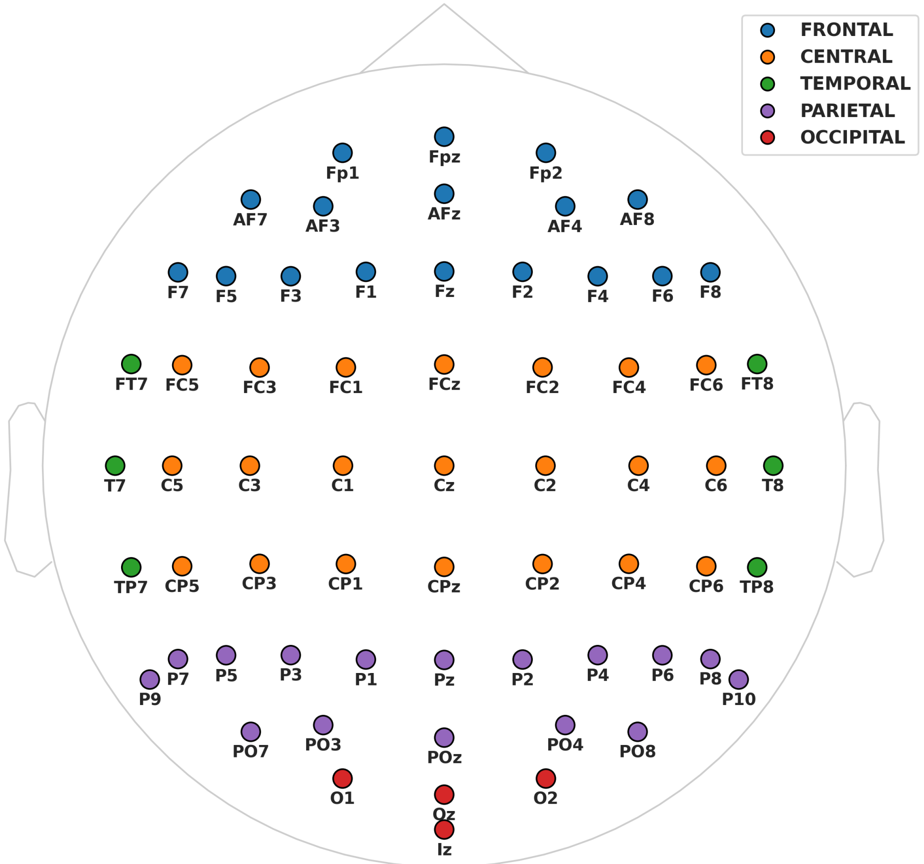


**Figure 8**

*Figure S8. BioSemi-64 electrode layout showing the scalp-region groupings used in regional analyses. Electrodes were assigned to five anatomical regions: frontal (blue), central (orange), temporal (green), parietal (purple), and occipital (red). Regional groupings were used to summarize encoding performance and characterize the spatial distribution of brain–language alignment across the scalp.*

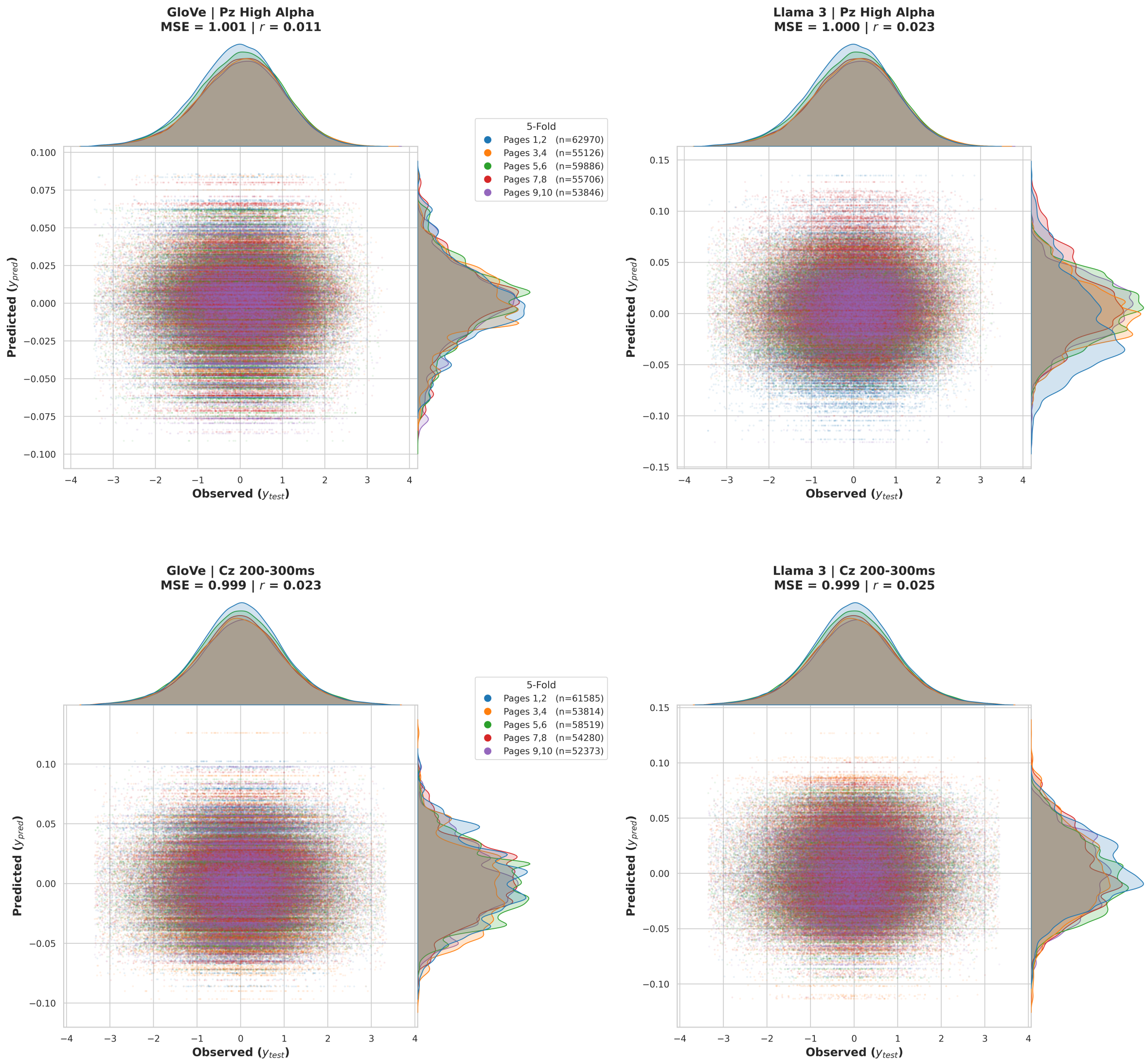


**Figure 9**

*Figure S9. Representative scatter plots comparing observed and predicted neural responses. Results are shown for PSD high-alpha activity at Pz and FRP activity during the 200-300 ms post-fixation interval at Cz using GloVe and Llama 3 embeddings. Each point corresponds to a fixation sample from the held-out test set, with colors indicating the five page-level cross-validation folds. Marginal density plots show the distributions of observed and predicted responses within each fold.*
